# When South meets North: a joint contact zone coinciding with environmental gradients in three boreal tree species

**DOI:** 10.64898/2026.03.13.711305

**Authors:** Pilar Herrera-Egoavil, J. Luis Leal, Qiujie Zhou, Pascal Milesi, Martin Lascoux, Burçin Yildirim

## Abstract

Post-glacial recolonization of Fennoscandia created secondary contact zones in many species, offering opportunities to study how gene flow and selection contribute to their establishment and maintenance. Here, we analyse genomic data from three boreal tree species—*Picea abies*, *Betula pendula*, and *Pinus sylvestris*—sampled along a latitudinal gradient in Sweden. Despite differences in colonization timing and dispersal ecology, all three species exhibit north–south genetic structuring aligned with environmental gradients. Most notably, the two main genetic clusters within each species overlap in a shared contact zone, corresponding to the climatic transition between Sweden’s two major environmental zones. The extent and structure of the contact zone differ among species: *P. abies* shows stronger genetic structure and moderate gene flow, *B. pendula* exhibits intermediate differentiation and gene flow, and *P. sylvestris* displays the weakest structure with stronger gene flow. All three species also show genomic signatures of local adaptation, with distinct underlying architectures. In *P. abies*, adaptive loci are broadly distributed across the genome, while, strikingly, they are mostly found within an inversion on chromosome 1 in *B. pendula*. In *P. sylvestris*, local adaptation likely relies on subtle allele frequency shifts across many loci with weak signals. These patterns align with theoretical expectations for polygenic local adaptation under varying migration regimes. Our comparative approach demonstrates how gene flow and selection jointly shape genomic landscapes in shared environments and contributes to understanding local adaptation in forest trees, with implications for predicting species’ responses to climate change.

## Introduction

Climatic oscillations during the Quaternary in Europe caused repeated expansions and contractions of species ranges. In particular, the Last Glacial Maximum (LGM, around 18,000 years ago) strongly shaped current species distributions across northern Europe (Petit et al., 2003). During this glacial period, species survived in more or less isolated areas called refugia, located south of the ice sheet (Hewitt, 1999). Some refugia were restricted to the southern peninsulas (Balkans, Italy, and the Iberian Peninsula), but fossil evidence indicates that many cold-adapted species also persisted at higher latitudes (Birks & Willis, 2008; Hošek et al., 2024). When the ice retreated and the climate became favorable, these lineages expanded their ranges, met again, and formed secondary contact zones (Petit et al., 2003). These zones are typically characterized by clines, spatial gradients, in phenotypic traits or allele frequencies that can persist despite ongoing gene flow (Hewitt, 1988; Kawakami & Butlin, 2012). The limits of the secondary contact zone and the degree of genetic differentiation across it would depend on several factors, including the duration of isolation, the effective population size of the two lineages, and migration rates (Ravinet et al., 2017).

Across Europe, many well-documented contact zones are the result of postglacial colonization of northern regions from southern refugia after the LGM, presenting a systematic distribution pattern rather than being randomly scattered over the continent (Schmitt, 2007). In Scandinavia, a recent contact zone emerged following the melting of the ice cap during the last stages of the LGM, enabling interaction between southwestern and northeastern genetic lineages. This contact zone in central Sweden is reported for different species, including *Clethrionomys glareolus* (Tegelström & Jaarola, 1998), *Ursus ursus* (Taberlet et al., 1995, Waits et al., 2000), *Picea abies* (Li et al., 2022; Nota et al., 2022), *Bufo bufo* (Thörn et al., 2021), *Phylloscopus sps.* (Hansson et al., 2000, Lundberg et al., 2023) and *Alnus glutinosa* (Havrdová et al., 2015). Such multispecies contact zones are often termed *suture zones*, referring to regions where the contact zones of different species geographically overlap (Johannesson et al., 2020; Remington, 1968). Similar patterns have been documented in other parts of the world, in both terrestrial (Hewitt, 1996; Swenson & Howard, 2005) and marine systems (DiBattista et al., 2015; Johannesson & André, 2006; Stanley et al., 2018). Comparative studies of these zones across species with different biological attributes could inform us about the factors shaping and maintaining barriers to gene flow. In particular, this would inform us on the impact of species-specific life history traits and genomic architecture (Chambers et al., 2023; Johannesson et al., 2020; Kawakami & Butlin, 2012).

Studying secondary contact zones in forest trees is particularly informative for understanding the strength of natural selection. Gene flow in temperate and boreal forest tree species is extensive, and occurs through both pollen and seed dispersal. Despite this extensive gene flow, they exhibit strong local adaptation and maintain high genetic differentiation (Savolainen et al., 2007; Kremer et al., 2012). Thus, in contact zones, genomes would be expected to homogenize rapidly under gene flow unless selection is strong enough to counteract it. Indeed, Li et al. (2022) recently demonstrated that *Picea abies* populations in Sweden consist of two major genetic clusters—a northern and a southern one—reflecting two recolonization routes after the LGM (Brewer et al. 2017), originating from populations with different genetic backgrounds (Zhou et al., 2024). They also demonstrated that the contact zone separating these clusters coincides with the transition between two main climatic zones in Sweden and is maintained by natural selection, with trees from each cluster adapted to their local environments.

Paleoecological studies further support that this contact zone formed as a result of recolonization of Scandinavia from both the south and the north. Pollen fossil data show that Norway spruce (*Picea abies*) first entered Scandinavia from the south around 12,000 years before present (YBP) and later from the north at about 10,500 YBP (Brewer et al., 2017; Giesecke, pers. comm.). Other dominant boreal forest tree species in Sweden, Scots pine (*Pinus sylvestris*) and birch (*Betula spp.*), also recolonized from both directions but somewhat earlier than *P. abies*. *P. sylvestris* arrived first from the south at ∼14,000 YBP and from the north at ∼10,000 YBP, while *Betula spp.* first appeared in the south at ∼14,000 YBP and in the north at ∼13,000 YBP (Brewer et al., 2017). Pollen abundance data further suggest that both *P. sylvestris* and *Betula* spp. were much more abundant than *P. abies* and recolonized Scandinavia more rapidly (Brewer et al., 2017).

While paleoecological evidence indicates that *Betula spp.* and *Pinus sylvestris* also recolonized Scandinavia from both southern and northern routes, it remains unclear whether a comparable contact zone persists and whether two distinct genetic clusters are present in these species, as observed in *P. abies*. The three focal species differ in life-history traits and ecological strategies, which may influence how contact zones are formed and maintained. *Betula pendula* is a monoecious, short-lived, wind-pollinated angiosperm tree species. Its distribution spans most of Europe and central Siberia, with a patchy presence in the south and a continuous range in northern regions. It is an important ecological component of boreal forests and a pioneer species that contributes to secondary vegetation succession (Beck et al., 2016). *Picea abies* and *Pinus sylvestris* are monoecious, wind-pollinated gymnosperms. *P. abies* occurs throughout Europe, mainly in mountainous areas of Central Europe and across Northern and Eastern Europe, extending to the Ural Mountains (Caudullo et al., 2016). It is a major component of boreal and subalpine conifer forests. Its eastern limit is not well defined due to extensive hybridisation with its close Siberian relative *Picea obovata*, with which it forms a syngameon (Zhou et al., 2024). In contrast, the range of *P. sylvestris* is continuous from Western Europe to Eastern Siberia, south to the Caucasus and Anatolia, and north to the Arctic Circle in Fennoscandia (Durrant et al., 2016). The two gymnosperm species differ in their colonization ability: *P. abies* tends to establish under tree cover, whereas *P. sylvestris* can recolonize open areas with poor soils (Mátyás, et al., 2004; Skrøppa, 2003). Overall, population genetic differentiation is low among the three species within their western ranges, though they differ in the degree of population structure they exhibit (Milesi et al., 2024). Population structure is pronounced in *Picea abies* (Tsuda et al., 2016; J. Chen et al., 2019; Zhou et al., 2024), moderate to weak in *Betula pendula* (Tsuda et al., 2017), and weak in *Pinus sylvestris* (Bruxaux et al., 2024; Kastally et al., 2025).

In this work, we leverage existing and newly acquired genomic data to study the three major boreal forest tree species: *Betula pendula* (silver birch), *Picea abies* (Norway spruce), and *Pinus sylvestris* (Scots pine). Specifically, we aim to (1) determine whether a contact zone is maintained in *B. pendula* and *P. sylvestris, separating genetically distinct northern and southern lineages* and, if so, identify its location for each species; (2) test for genomic signatures of selection potentially involved in maintaining the contact zone; and (3) compare the adaptive architectures among the three species.

## Material and Methods

### 1. Sampling and genotyping

For the three species, we assembled genomic datasets for trees sampled from populations distributed along a similar latitudinal gradient across Sweden (Figure 1A). For *Betula pendula*, we used exome capture data from 150 trees sampled in 12 Swedish populations (see details in Supplementary Methods and Leal et al. (2024)). For *Picea abies*, raw reads from exome capture sequencing were downloaded from NCBI (Bioproject PRJNA511374, Chen et al., 2019, and Bioproject PRJNA731384, Z.-Q. Chen et al., 2021) for 154 trees, pooled into 12 populations, for which sampling location and genotype assignments confirmed Swedish origin (Li et al., 2022). For *P. sylvestris*, raw reads from genotyping-by-sequencing (GBS) were downloaded from NCBI (accession number PRJNA976641, Bruxaux et al., 2024) for 351 trees from 10 Swedish populations.

**Figure 1:**
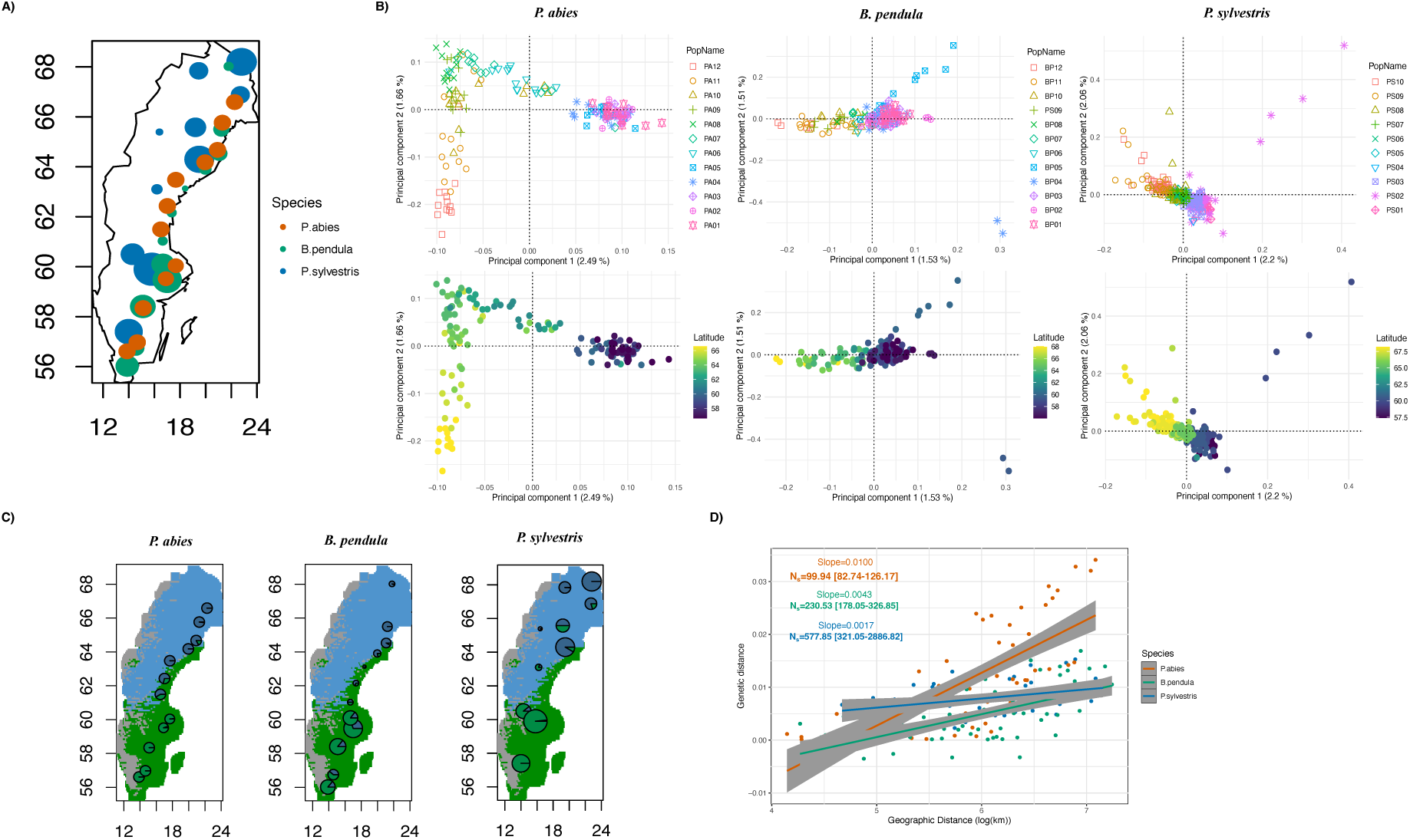
Samples and population structure of three species A) Sampling locations for three species. The size of the dots represents the sample size for each population B) The first two principal components from PCA, plotted by population (ordered from north to south in the legend) and latitude C) Ancestry proportions for each population based on the ADMIXTURE model (K = 2), with colors representing different ancestry components. The background colors corresponds to the climatic zones, climatic zone 1 (blue), climatic zone 2 (green), climatic zone 3 (grey). D) Isolation by distance (IBD) patterns for each species, along with the linear regression results and neighborhood size (N_S_) estimates.

Raw reads were mapped with BWA-MEM (v.0.7.17, Li, 2013) against the respective reference genome for *B. pendula* (Bpev01; Salojärvi et al., 2017) and *P. abies* (Pabies1.0; Nystedt et al., 2013), and the phylogenetically closest reference for *P. sylvestris* (*Pinus tabuliformis*, P.tabuliformis_V1.0; Niu et al., 2022; Kesälahti et al., 2025). SNP calling and filtering were then performed using GATK (v.4.3.0.0; DePristo et al., 2011). For all three species, SNPs with more than 50% missing data were excluded using VCFtools (v.0.1.16, Danecek et al., 2011), along with SNPs overlapping with putative multi-copy regions identified with the R package rCNV (v.1.3.0, Karunarathne et al., 2023). Due to differences in sequencing technologies and reference genome quality, species-specific filtering criteria and parameters were applied where relevant (see Supplementary Methods for details).

The final datasets contained 597,182, 51,768 and 86,722 SNPs for *P. abies*, *B. pendula* and *P. sylvestris*, respectively. The SNPs were functionally annotated based on the annotation available for the respective reference genomes and classified as intergenic, intronic, synonymous and nonsynonymous using SNPEff (v.4.3t; Cingolani et al., 2012) for *P. abies* and *B. pendula*, and ANNOVAR (Wang et al., 2010) for *P. sylvestris*. Intergenic, intronic and synonymous sites were considered putatively neutral, resulting in 415,216, 23,684 and 66,196 SNP datasets for *P. abies*, *B. pendula* and *P. sylvestris*, respectively.

In both *P. abies* (genome size ∼20 Gb) and *B. pendula* (genome size ∼440 Mb), SNPs were obtained through exome capture, whereas in *P. sylvestris* (genome size ∼22 Gb), they were generated using genotyping-by-sequencing (GBS). The GBS approach used for *P. sylvestris* might be expected to capture less genes than exome capture. However, the proportion of SNPs located in genic regions (intron+synonymous+nonsynonymous) was comparable across datasets, despite the different sequencing approaches (Supplementary Methods). We therefore believe that the use of GBS versus exome capture does not introduce substantial bias in our comparisons.

### 2. Population structure and spatial pattern of variation

To investigate population structure in each species we first performed principal component analysis (PCA) using PLINK (v.1.9., Chang et al., 2015). We then inferred ancestral components using ADMIXTURE (v.1.3, Alexander et al., 2009) and TESS3 (v.1.1.0, Caye et al., 2016), representing non-spatial and spatially explicit models, respectively. These analyses were performed using neutral SNP datasets after removing singletons and variants in high linkage disequilibrium (*r*^2^ > 0.2) using PLINK (Chang et al., 2015).

We quantified the intensity of isolation-by-distance (IBD) by regressing genetic distance (F_ST_/(1-F_ST_)) against the logarithm of geographic distance (Wright, 1943; Rousset, 1997). Weir and Cockerham’s population pairwise F_ST_ values were calculated from neutral SNPs using VCFTools (Danecek et al., 2011; Weir & Cockerham, 1984) and geographic distance (geodesic) with the R package geosphere (v.1.5-18, Hijmans, R.J., 2022). In continuously distributed populations, the inverse of the regression slope between genetic and geographic distance is expected to be proportional to the neighborhood size (N_S_), which describes the dispersal potential of individuals within a population (Rousset, 1997). As the species we examined are continuously distributed across Sweden, we interpreted the slope as an approximation of N_S._ Next, we estimated effective migration surfaces across the species’ ranges using FEEMS (v.1.0.1, Marcus et al., 2021), after removing SNPs with a minor allele frequency of less than 1% from the full datasets; datasets including only putatively neutral SNPs gave the same results.

Finally, we assessed environmental variation across the sampling range. Briefly, we obtained 19 bioclimatic variables from the CHELSA database (v.1.2, https://chelsa-climate.org/; Karger et al., 2017) across Sweden. The first 11 variables are temperature-related, and the remaining variables are related to precipitation. Based on these data, we performed hierarchical clustering to define major climatic zones using the FactoMineR package in R (v.2.11, Lê et al., 2008), following Li et al. (2022).

### 3. Barriers to gene flow

To examine the potential contribution of selection to the genetic divergence between northern and southern genetic clusters, we used the program RIDGE (v.1, Burban et al., 2024). RIDGE employs an Approximate Bayesian Computation (ABC) approach to classify loci as barriers between two diverging lineages. The program considers four demographic models: Strict Isolation (SI), Isolation with Migration (IM), Secondary Contact (SC), and Ancestral Migration (AM), alongside four genomic models that incorporate either homogeneous or heterogeneous effective population size (N_e_) due to linked selection (1N, 2N) and homogeneous or heterogeneous migration rates due to selection at barrier loci (1M, 2M). Given that migration is not involved in the Strict Isolation model, this results in 14 demographic × genomic models. Instead of inferring model parameters for a single best model, RIDGE analyzes all models and assigns weights based on their fit to the observed data, creating a hypermodel. Under this hypermodel, a posterior distribution of each parameter is obtained, and a goodness-of-fit value is calculated. This approach allows the classification of loci as barrier or non-barrier without introducing bias due to model selection.

We ran RIDGE on the full datasets for two genetic clusters of each species, using a 10 kb window size for *P. abies* and *B. pendula*, and a 100 kb window size for *P. sylvestris*, based on the SNP density in our datasets (details of all runs are provided in the Supplementary Methods). To evaluate whether genetic distance between individuals from the northern and southern clusters influences RIDGE results, we repeated our analyses for each species using three different datasets, based on genetic distance. The ancestral coefficients for the first ancestral component derived from ADMIXTURE for K=2, corresponding to the northern clusters, were used to define genetic similarities (Li et al., 2022). Individuals with ancestral coefficients between 0.4 and 0.6 were categorized into the ‘close’ datasets. Those with coefficients between 0.1 and 0.4 or between 0.6 and 0.9 were included in the ‘intermediate’ datasets. Finally, individuals with ancestral coefficients below 0.1 or above 0.9 were classified into the ‘far’ datasets. For each species, the datasets contained 10 randomly sampled individuals, five from each genetic cluster. RIDGE was run separately for each dataset, and the results were compared with the full dataset.

### 4. Signatures of local adaptation

To investigate and compare the role of adaptation in shaping genetic diversity along latitude in the three species, we conducted a series of genome scans and genotype-environment association (GEA) studies (correlations between allele frequencies and environmental variables with Bayenv2). Two genome scan approaches were used to identify loci showing extreme differentiation relative to the genomic background. The first, *pcadapt*, is an individual-based approach that does not require predefined population assignments (*pcadapt* v.4.4.0 R package; Luu et al., 2017; Privé et al., 2020). It performs principal component analysis on the centered and scaled genotype matrix and assumes that most SNP variation reflects population structure. SNPs showing stronger differentiation than expected under this structure are identified as outliers and considered candidates for selection. For all species, we used the first two principal components to account for population structure in this analysis (see Supplementary Methods). For the second genome scan, we used a population-based approach, X^T^X statistic from Bayenv2 (Coop et al., 2010; Günther & Coop, 2013). X^T^X is an F_ST_-like measure based on standardized allele frequencies that are corrected for population structure. Extreme values therefore reflect differentiation presumably caused by selection. Population structure was accounted for by estimating the variance–covariance matrix among populations from putatively neutral SNPs using Baypass (v.2.1; Gautier, 2015). Finally, for each of the 19 bioclimatic variables mentioned above, we conducted genotype–environment associations (GEA) using Bayenv2 (Coop et al., 2010; Günther & Coop, 2013).

These analyses were conducted on the full SNP datasets pruned for SNPs with a minor allele count < 10 for *B. pendula* and < 15 for *Picea abies* and *Pinus sylvestris*, keeping 95,680, 16,794 and 20,787 SNPs for *P. abies, B. pendula* and *P. sylvestris*, respectively. For each analysis, we applied stringent criteria to control for false-positives detection risk (see Supplementary Methods).

### 5. Analysis of sharp change in allele frequency across the contact zone

Spatial gradients in allele frequencies across contact zones are referred to as clines. To test for sharp allele frequency transitions and estimate selection strength at loci involved in the maintenance of the contact zone, we fitted explicit cline models (Szymura & Barton, 1986, 1991) to each SNP using the R package hzar (v.0.2.5, Derryberry et al., 2014) in order to estimate their center (*c*) and width (*w*). The cline center represents the geographic location where allele frequency changes are steepest. The width, defined as the inverse of the maximum slope at this center, is proportional to the ratio of dispersal to the square root of selection strength 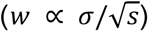 (Barton & Gale, 1993) with *σ* being the standard deviation of the distance between parent and offspring measured along a linear axis and *s* the selection intensity. Thus, with dispersal estimates, the width parameter allows for calculation of selection intensity.

Cline fitting was performed using all SNPs in the final datasets. Allele frequencies per population were used as the response variable, and geographic distance from the northernmost site in Sweden (69.06°N, 22.55°E) was the predictor. We fitted four models differing in shape (sigmoid or stepped clines) and the number of estimated parameters (Szymura & Barton, 1986). Model I estimated only the cline center (*c*) and width (*w*), fixing allele frequencies at range edges *(p*_*min*_, *p*_*max*_), thus producing a sigmoid cline. Model II also estimated *p*_*min*_ *and p*_*max*_, allowing for more flexible sigmoid shapes. Model III and Model IV were stepped versions of Models I and II, incorporating exponential tails flanking the central cline and requiring four additional parameters to define the left and right tails (Kawakami & Butlin, 2012). The model with the lowest Akaike’s information criterion, corrected for small sample sizes (AICc), was defined as the best-fitting model. We then used the AICc weight of the best-fitting model and of a null model representing a uniform distribution. The ratio of these AICc weights, termed evidence ratios, quantifies how much more likely one model is compared to the other (Anderson, 2008), *i.e.,* cline vs null model. Instead of defining an arbitrary threshold, we adopted an outlier approach: an allele was classified as clinal if its evidence ratio for the best-fitting cline model, relative to the null model, exceeded the 99th percentile of the distribution across all loci. Hereafter, we refer to alleles satisfying this criterion as clinal alleles.

Note that, if allele frequencies change linearly across the geographic range, the model either fails to converge or estimates very large widths (approaching infinity), indicating the absence of a sharp transition zone and cline center. Such loci are not considered as true clines, and our method inherently excludes linear gradients.

We next tested whether SNPs identified in genome scans and GEA analyses (selection candidates) were enriched for clinal alleles using a hypergeometric test, comparing them to all SNPs with similar allele frequency differences between cline ends. For these presumably selected and clinal alleles, we examined the distributions of center (*c*) and width (*w*) values. Additionally, to estimate selection strength at these loci, we used neighborhood size, previously estimated from the isolation-by-distance analysis based on neutral SNPs,as a proxy for dispersal rate, N_S_ ≈ *σ*² (Slatkin & Barton, 1989; Rousset, 1997). Selection strength was then calculated from cline width as *s* ≈ N_S_/*w²*.

Finally, to assess whether environmental variables themselves exhibit clinal transitions across the contact zone and whether these align spatially with allele frequency clines, we applied the same cline-fitting procedure to each of the 19 bioclimatic variables and to the first principal component (PC1) of a PCA conducted on them, using normalized bioclimatic values ([0–1]) at each population location as the dependent variable.

### 6. Functional annotation

For all species, we extracted coding sequences (CDS) of genes annotated in the reference genomes and translated them into protein sequences using *gffread* (v.0.12.7, Pertea & Pertea, 2020). Each protein sequence was aligned against two databases - RefSeq non-redundant and UniProtKB—using DIAMOND (v.2.1.9, Buchfink et al., 2021). We retrieved the five most significant hits from each alignment and obtained Gene Ontology (GO) terms (biological process only) through ID mapping (https://www.uniprot.org/id-mapping). We also obtained GO terms by scanning protein sequences with InterProScan (v.5.62-94.0, Jones et al., 2014). Functional enrichment analyses of the GO terms were performed with the topGo R package (v.2.58.0, Alexa et al., 2006) against a background set that included genes linked to SNPs in our initial vcf file, to avoid potential bias from the reduced representation of the genomes due to the sequencing techniques. Enriched GO terms were summarised with rrvgo (v.1.18.0, Sayols, 2023) and visualized with ggplot R package (v.3.5.2, Wickham, 2016).

## Results

### 1. The contact zone is retrieved in the three species

The first two principal components (PC) explained approximately 3-4% of the genetic variation in all three species (Figure 1B). For all three species, the first PC separated the northernmost and southernmost populations, with a gradient between them. The second PC for *P. abies* further differentiated the northern populations, while no distinct pattern was observed for *B. pendula* and *P. sylvestris*.

Cross-validation curves from ADMIXTURE did not plateau for any of the species (Figure S1A). TESS3 cross-validation scores likewise showed no plateau but showed increasing support for K from 1 to 3 (Figure S1B). This indicates no strong support for a specific K. In such cases, caution should be given when interpreting K values, as the number of genetic groups detected does not necessarily correspond to the number of biologically meaningful populations in the sample (François & Durand, 2010). Moreover, the estimation of K strongly depends on sampling, and we lack populations outside Sweden. However, since previous studies including broader sampling in *P. abies* and *P. sylvestris* reported two ancestry components (Li et al., 2022; Hall et al., 2021), and we observed latitudinal genetic differentiation in all species, we examined ancestry proportions for K=2. At this K, ADMIXTURE separated populations in the northern and southern regions of Sweden into two genetic clusters (Figure 1C), a pattern also recovered by TESS3 (Figure S1C).

The contact zone between the northern and southern genetic clusters, where the most admixed individuals were found, was geographically similar across all three species, spanning latitudes from 60°N to 63°N. In *P. abies*, the southern ancestry component contributed to individual genetic diversity at latitudes up to 65°N, yet overall, genetic exchange between northern and southern clusters outside the contact zone was more limited compared to the other two species. In *P. sylvestris*, southern ancestry component contributions extended up to latitude 67°N, whereas in *B. pendula*, the northern ancestry component contributed more extensively to southern populations beyond the contact zone.

The overall pairwise F_ST_ values across species sampling ranges were low — averaging 0.0121 (sd=0.0094), 0.0049 (sd=0.0049), and 0.0077 (sd=0.0032) for *P. abies*, *B. pendula*, and *P. sylvestris*, respectively (Figure S2A). Within species, differentiation was slightly higher between northern and southern populations, being most pronounced in *P. abies* and least in *P. sylvestris* (Figure S2B), consistent with a decreasing population structure from *P. abies* to *P. sylvestris*. All three species exhibited significant isolation-by-distance (IBD) patterns, with the strongest in *P. abies* and the weakest in *P. sylvestris* (Mantel test on geographic and genetic distance matrices; *ρ*= 0.805, *P*-value=1×10^−4^, *ρ*= 0.682, *P*-value=1×10^−4^, *ρ*= 0.331, *P*-value=0.023, for *P. abies*, *B. pendula*, and *P. sylvestris*, respectively). Accordingly, neighborhood size increased from *P. abies* to *P. sylvestris* (Figure 1D). IBD patterns within northern and southern populations remained significant only for *P. abies* (for northern populations: *ρ*= 0.689, *P*-value=0.0097; for southern populations: *ρ*= 0.611, *P*-value=0.0056), but not for *B. pendula* or *P. sylvestris*. This suggests that in *B. pendula* and *P. sylvestris*, significant IBD patterns are driven primarily by comparisons between northern and southern populations, whereas in *P. abies*, genetic clusters contribute differently to the observed variation. Indeed, in *P. abies*, neighborhood size was larger within southern populations and smallest between northern and southern populations (Figure S3).

The effective migration surfaces estimated by FEEMS mostly corroborated the IBD patterns. For *P. abies*, we observed reduced migration rates within both the contact zone and the northern cluster. In contrast, for *B. pendula*, reduced migration rates were only observed around the contact zone (Figure 2). Compared to the other two species, migration rates in *P. sylvestris* were relatively homogeneous with the exception of the far northern region, where migration rates appear lower but this border effect could also be a consequence of sampling and should be interpreted carefully.

**Figure 2:**
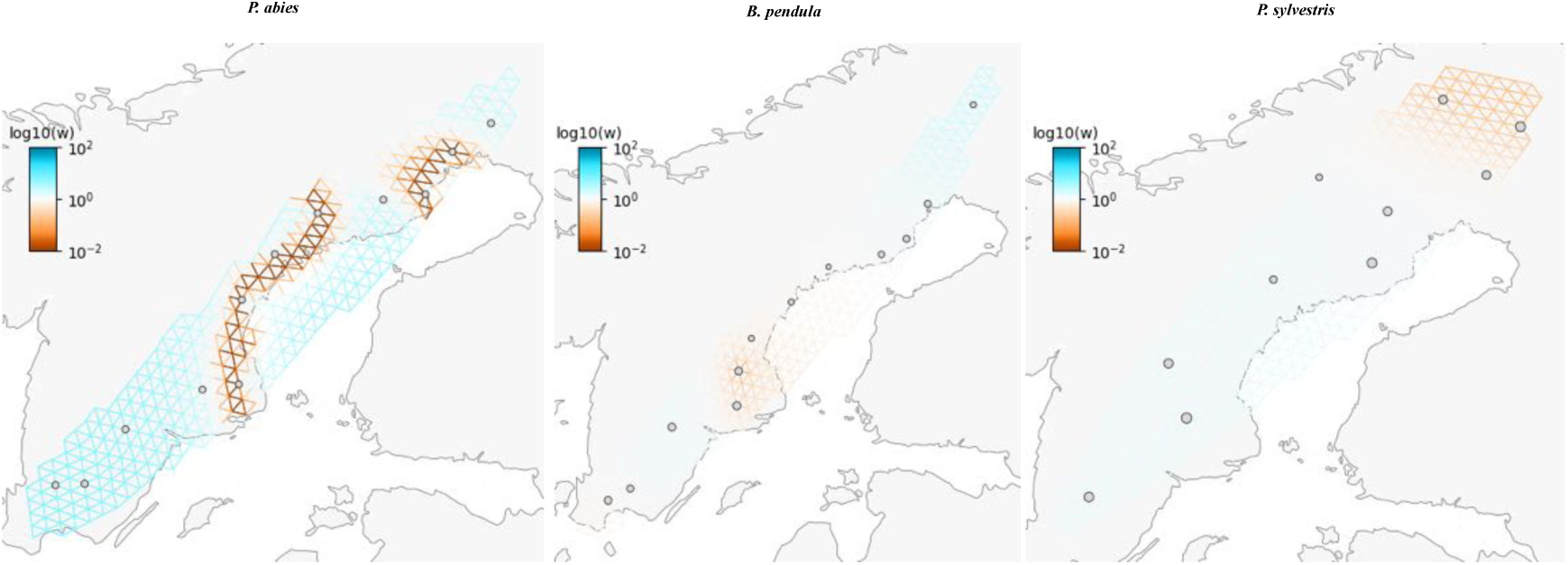
Effective migration rate surfaces across the distribution ranges of each species estimated using FEEMS. Blue and red colors indicate higher and lower migration rates, respectively, compared to the expected rates under a homogeneous IBD model.

To investigate whether the contact zone between the two main genetic clusters corresponds to transition in the environment, we identified three climatic zones using an unsupervised clustering approach with 19 bioclimatic variables. The first two environmental clusters separated the northern and southern parts of Sweden, and were primarily based on temperature-related variables (Figure S4). The third cluster, located in the western part of the country, was distinguished by precipitation-related variables. The transition between the northern and southern environmental clusters overlapped with the genetic contact zone observed in all species (Figure 1B). Using the first ancestral component from the ADMIXTURE analysis with K = 2, which corresponds to the northern genetic cluster, as a reference, we analyzed the distribution of ancestral coefficients across individuals in each climatic zone. The coefficients differed significantly between climatic zones, indicating that trees from the northern and southern genetic clusters originate predominantly from their respective climatic zones (Figure 3). In contrast, highly admixed individuals, with ancestral coefficients between 0.3 and 0.6, were more evenly distributed across both climatic zones. Despite the similar distributions of highly admixed individuals across climatic zones, the differences in ancestral coefficients for these individuals were low but significant for *P. sylvestris* and *B. pendula*.

**Figure 3:**
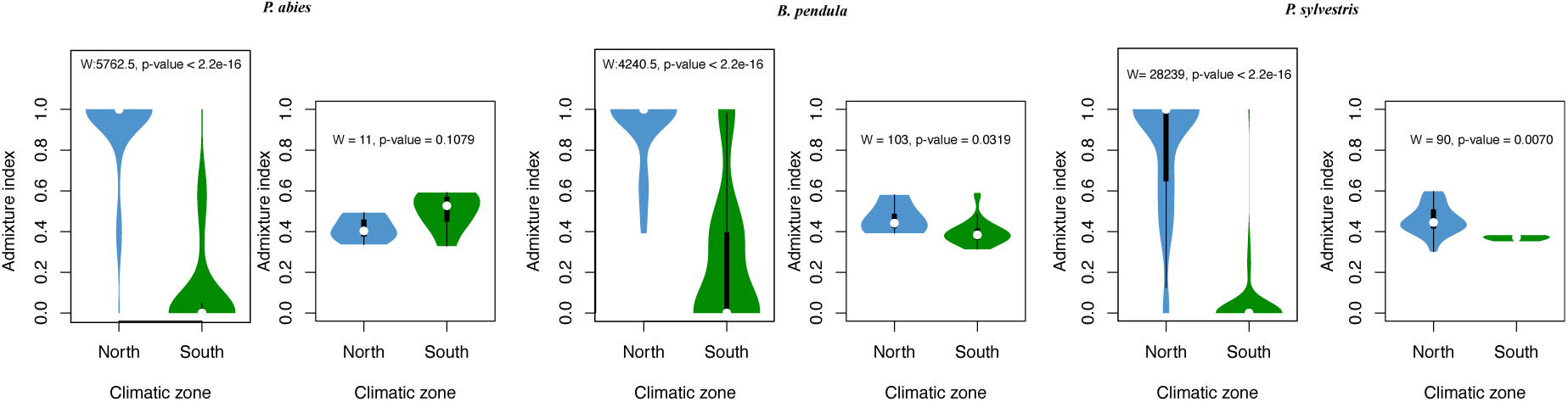
Distribution of the admixture index for the first ancestral component from ADMIXTURE analyses across each climatic zone. A higher admixture index corresponds to a greater northern genetic ancestry. For each species, the left panels display the distribution for all individuals, while the right panels show the distribution for highly admixed individuals, with admixture indices between 0.3 and 0.6. The text in the plots presents the results of the Wilcoxon rank-sum test between the climatic zones.

### 2. Selection contributes to the contact zone

We investigated whether selection contributes to maintaining the genetic divergence between the northern and southern clusters of each species by creating barriers to gene flow, using RIDGE software. Our analyses on the full dataset (between northern and southern genetic clusters) successfully fitted the parameters to the observed data, with goodness-of-fit values for each species exceeding the rejection threshold of 0.05 (0.316, 0.633, and 0.618 for *P. abies*, *B. pendula*, and *P. sylvestris*, respectively). In all three species, the model weights indicated a preference for demographic models involving ongoing migration, which are isolation with migration (IM) and secondary contact (SC), over ancestral migration (AM) and strict isolation (SI) (Figure 4). Additionally, the genomic models supported a heterogeneous landscape along the genome for both migration and effective population size. These results confirm that current gene flow occurs in each species and suggest that selection also influences the observed genetic divergence. The analysis performed on the datasets separated by the genetic distance (close, intermediate and far) supported the same hypermodel with ongoing migration and heterogeneous genomic landscape caused by linked selection (Figure S5).

**Figure 4:**
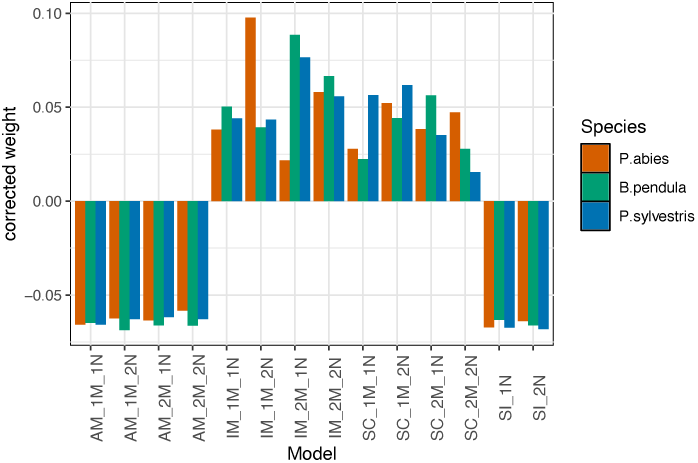
Model weight of 14 models for each species corrected by the uniform distribution model weight.

### 3. Different genetic architectures and degrees of local adaptation across species

We tested for the presence of loci contributing to local adaptation through genome scans (*pcadapt* and X^T^X statistics-Bayenv2) and genotype-environment association (GEA) analyses (Bayenv2). For *P. abies*, we identified 39 and 806 SNPs showing extreme allele frequency differentiation, using *pcadapt* and X^T^X, respectively. For *P. sylvestris*, we detected 42 candidate SNPs using *pcadapt* and 92 using X^T^X. In both species, the candidates were distributed across the genome (Figure 5A, Table S1). While in *B. pendula*, 119 (*pcadapt*) and 137 (X^T^X) SNPs identified as candidates were mostly clustered in chromosome 1 between positions 33,516,284 and 42,732,563 (∼9 Mbps). The loadings from *pcadapt* showed that variants within this region strongly contributed to the first two principal components (Figure S6A). To assess whether this region was masking additional signals, we performed a second genome scan after excluding it. However, no additional candidate SNP were detected (Figure S6B). To investigate whether this genomic region harbors an inversion, we calculated pairwise LD across all SNPs on chromosome 1. The resulting LD heatmap revealed a distinct long-range linkage pattern within this region, which was absent in the rest of the chromosome and in other chromosomes (Figure S7), supporting the presence of an inversion.

**Figure 5:**
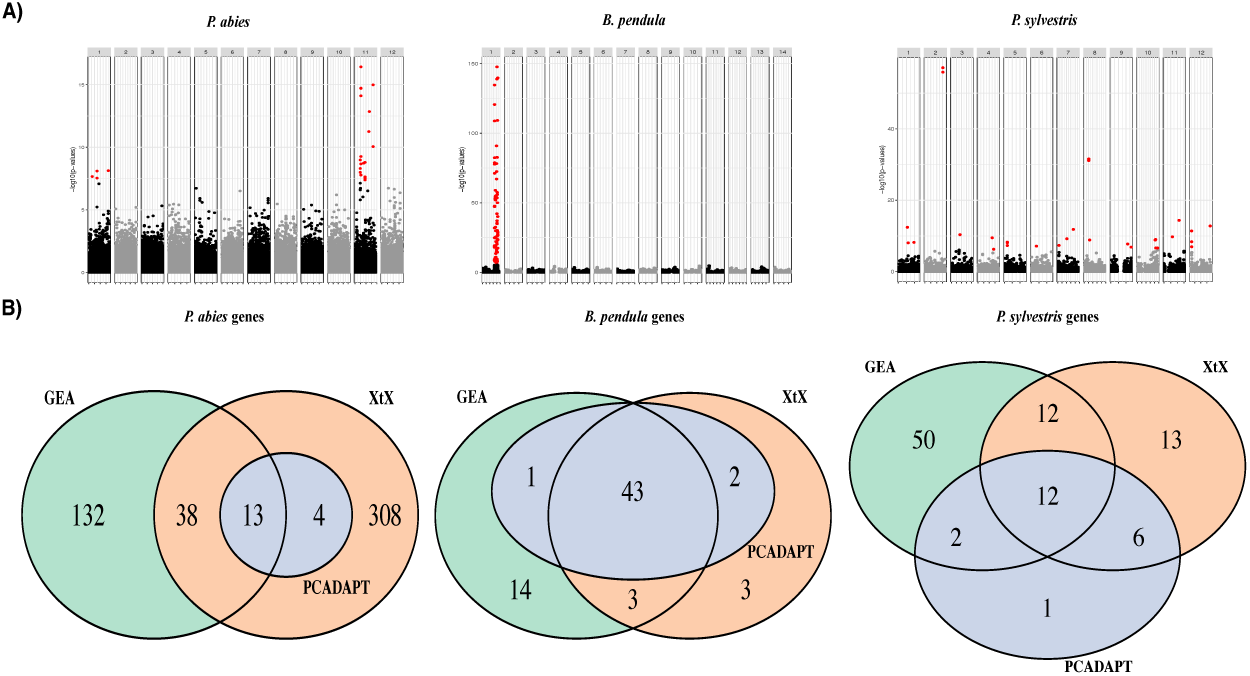
Candidate SNPs and genes identified in genome scans and genotype-environment association (GEA) analyses A) Manhattan plots of genome scan for an excess of differentiation (*pcadapt*), with candidate SNPs marked in red. B) Number of overlapping and unique genes identified in different genome scans (*pcadapt*, X^T^X) and GEA analyses.

The GEA analyses identified 300 (*P. abies*), 142 (*B. pendula*) and 142 (*P. sylvestris*) candidate SNPs associated with at least one of the 19 environmental variables (Table S1). Candidate SNPs were distributed across the genome for *P. abies* and *P. sylvestris*, while for *B. pendula*, most candidate SNPs were on chromosome 1, within the putative inversion. At the gene level, there were varying levels of overlap across different tests within each species (Figure 5B). Across all analyses, *P. abies* showed a 76% overlap in candidate genes, *B. pendula* exhibited a higher overlap of 93% and *P. sylvestris* had 57% (Table S2). All in all, the proportion of adaptive SNPs and genes across the full dataset was similar for *P. abies* and *P. sylvestris* (0.17% and 0.22% for adaptive SNPs, and 1.77% and 1.65% for adaptive genes, respectively), while it was about twice as much for *B. pendula* (0.31% and 3.05%; Table S1). As across all analyses in *B. pendula*, at least 85% of adaptive variants come from the putative inversion region (100%, 91.2% and 85.2% for *pcadapt*, X^T^X and GEA, respectively), this elevated signal may reflect the inversion also capturing highly linked non-causal variants. Additionally, most candidate SNPs detected with genome scans were also identified with GEA in the three species except for X^T^X in *P. abies* where additional candidate genes were identified (Table S1, S2).

From the unique Gene Ontology (GO) terms for all candidate genes, 83 were common to all species (Figure S8A), but this overlap was not significantly higher than expected if GO terms were randomly selected from background genes (Table S3). Interestingly, these 83 GO terms were associated with functions such as biotic and abiotic responses, immune system processes, growth and development, and circadian clock regulation (Figure S8B). Enrichment analyses of candidate genes from all analyses resulted in 131, 14, and 25 significantly enriched GO terms (P-value < 0.01) for *P. abies*, *B. pendula*, and *P. sylvestris*, respectively (Figure S9). No GO terms were commonly enriched across all three species. Nevertheless, for all three species, the enriched GO terms were linked to biological processes such as flowering and vegetative timing, growth and development, photoperiodism, circadian rhythm, and salicylic acid signalling, all of which are known to be involved in plant responses to climatic stress (Gray & Brady, 2016; Whiting et al., 2024).

### 4. Allele frequency changes reflect environmental gradients across loci and species

To estimate the strength of selection at loci involved in maintaining the contact zone, we first fitted sigmoidal or stepped cline models to allele frequency changes across populations and estimated cline center and width for alleles showing clinal variation. In all species, clinal variation was best explained by a sigmoid function (Model I and II; Figure S10A). Thereafter, we refer to alleles fitting this sigmoidal pattern as *clinal alleles*. To assess fit quality, we calculated residual standard errors (RSE) between observed and fitted allele frequencies. RSE values were more variable and skewed toward higher values in *P. sylvestris* compared to the other species (Figure S10B), and *B. pendula* had slightly but higher RSE variance than *P. abies*.

SNPs identified by genome scans (*pcadapt* and X^T^X) were significantly enriched for clinal alleles in all three species (Table S4). In *P. abies*, SNPs identified in GEA analyses showed significant enrichment for clinal alleles in 9 out of 11 temperature-related variables, but in none of the precipitation-related variables. *B. pendula* showed a similar pattern, with enrichment in 10 temperature variables and none in precipitation variables. In contrast, *P. sylvestris* showed enrichment for clinal alleles across all environmental variables except one.

We also compared the prevalence of clinal alleles in neutral datasets. Across all species, clinal alleles were much less frequent among neutral SNPs than among putatively selected ones, and enrichment was not significant for *P. abies* or *B. pendula*, but was significant for *P. sylvestris* (Table S4). Still, selection candidates showed significantly higher AIC differences between the cline and null models than neutral SNPs, indicating stronger support for clinal patterns among selection candidates (Figure S10D).

#### Cline patterns and geographic positions of centers

In *P. abies*, all top candidate SNPs identified in genome scans and GEA analyses exhibited clinal patterns (Figure 6A). These SNPs were broadly distributed across the genome and showed variation in cline parameters (Figure 5A). Although the parameters formed a continuum, two main groups emerged with a smooth transition between them: one centered around 350 km south of the northernmost point of Sweden with narrower widths, and a second centered around 650 km with broader widths. The distribution of parameters for all putatively selected clinal SNPs mirrored those of the top SNPs (Figure 6B). The relatively southern center (∼650 km) corresponds with the contact zone and environmental transitions identified at latitudes 60–63°N, located 600–1000 km south of the northernmost point of Sweden (Figure 1C), and aligns with cline centers estimated for bioclimatic variables in *P. abies*, underscoring the dominant role of environmental selection (Figure S11).

**Figure 6:**
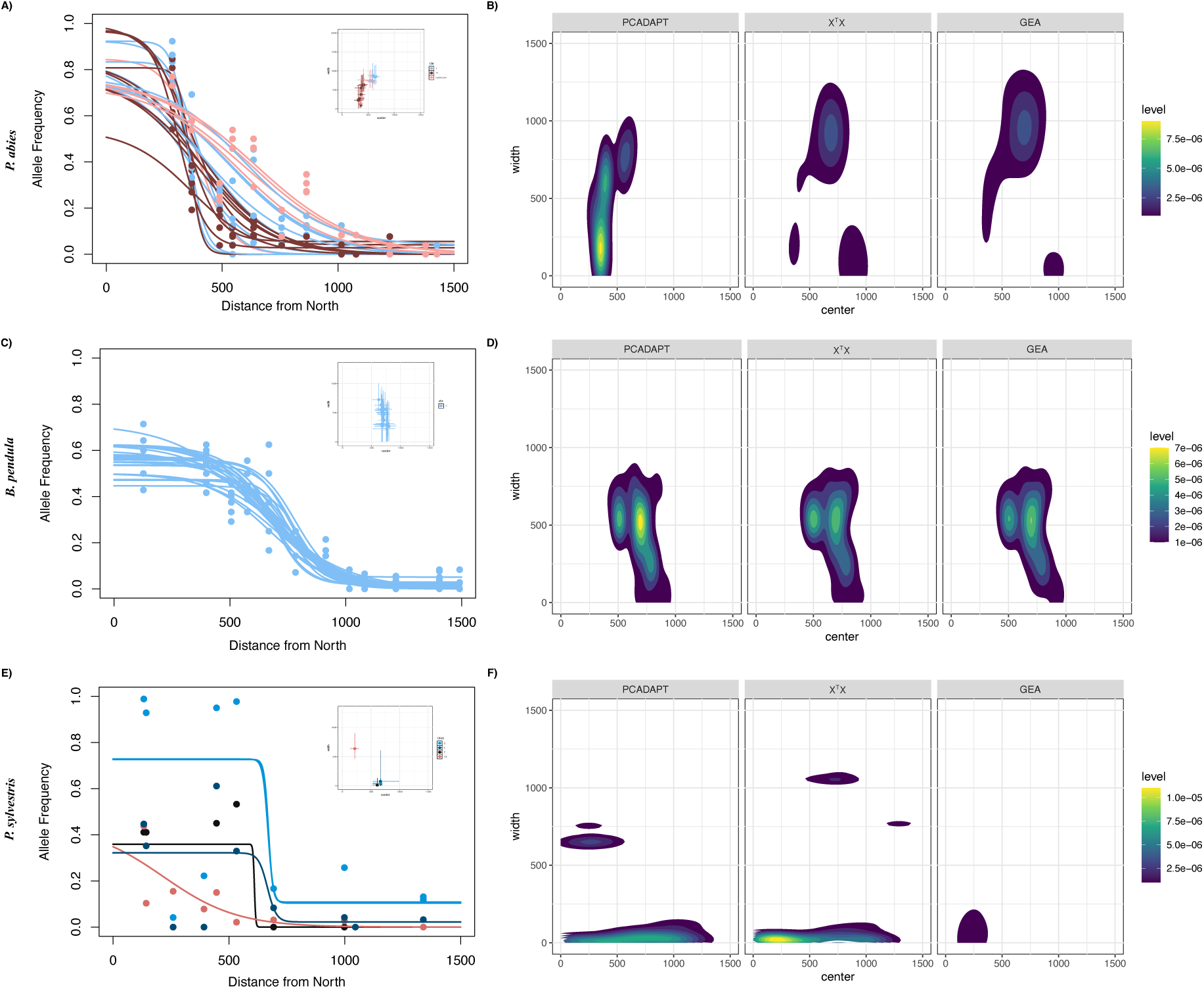
A) *P. abies* geographic clines for allele frequencies of top 25 SNPs identified commonly between genome scans and GEA analyses for climatic variable 1. The inset plot shows the parameter distribution (with 95% confidence intervals) for the same SNPs and colors represents SNPs from different chromosomes. B) Two dimensional density plots illustrating the distribution of center and width parameters for all clinal SNPs detected in genome scans and GEA analyses. Panels C (30 SNPs), D: same information for *B. pendula* and Panel E (5 SNPs), F: same information for *P. sylvestris*.

In *B. pendula*, most top candidate SNPs from genome scans and GEA analyses were located within the putative inversion on chromosome 1 and displayed strong clinal patterns with consistent centers and widths (Figures 6C). The distribution of parameters for all clinal SNPs resembled those of the top candidates (Figure 6D). Although a few clinal SNPs were found outside chromosome 1, they showed similar center positions and widths, suggesting a uniform clinal trend. Cline centers were concentrated between 500 and 1000 km, again overlapping with the contact zone and environmental transition (Figure 1C). In contrast, clinal patterns in *P. sylvestris* were more variable, consistent with higher RSE values. While some SNPs identified in selection scans were classified as clinal, only the top five—consistently detected across selection scans on different chromosomes—showed clear sigmoidal clines (Figure 6E). These SNPs had center estimates at ∼300 km and ∼650 km, with the latter matching the environmental transition and the former overlapping some bioclimatic variables (Figure S11). However, the overall distribution of clinal SNP parameters did not show clustering around a specific value (Figure 6F).

We next calculated selection strength for clinal and putatively selected alleles, detected in the previous steps (*pcadapt*, X^T^X, GEA), using estimates of neighborhood size and cline width (*s* ≈ N_S_/*w²*). The first observation for all species was the bimodal distribution of selection coefficients (Figure 7). This bimodality resulted from differences in width estimates between cline models: Model I produced higher width values and, consequently, lower selection coefficients compared to Model II (Figure S12B). To avoid inconsistencies, we did not compare estimates across different models but instead focused on comparing parameters within the same model across species.

**Figure 7:**
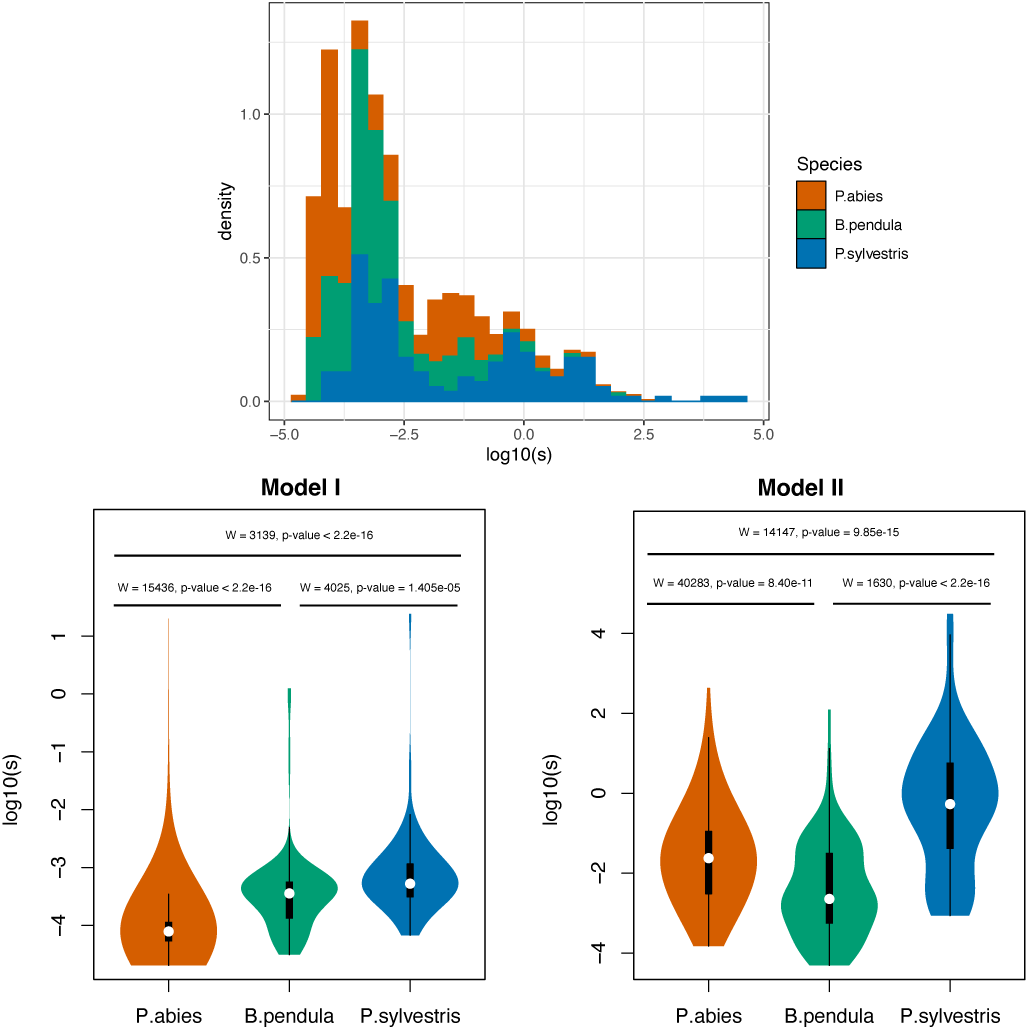
Selection coefficients calculated from cline widths for all putatively selected alleles. The upper panel shows the distribution of selection coefficients for all alleles in each species, while the lower panel presents the coefficients for alleles according to their best-fitted cline model.

Selection coefficients from Model I were highest in *P. sylvestris* and lowest in *P. abies*, with *B. pendula* showing intermediate values (Figure 7). In Model II, selection coefficients remained highest in *P. sylvestris*; however, *P. abies* exhibited higher values than *B. pendula*. Additionally, the qualitative pattern remained consistent when selection coefficients were examined separately for alleles detected in different selection scans (Figure S13).

## Discussion

### 1. A shared contact zone with varying levels of population structure

Our analyses identified two main genetic clusters within each species, which overlapped within a shared contact zone between 60° and 63° latitude north in Sweden. This contact zone also aligned with the transition between distinct climatic zones, indicating that environmental factors may play a role in maintaining genetic divergence. The same contact zone has been observed before for *P. abies* (Li et al., 2022) but not for *B. pendula* and *P. sylvestris*. As Li et al. (2022) also included populations outside Sweden and considered a larger number of individuals within Sweden, they had more power in differentiating the northern and southern genetic clusters via ADMIXTURE and in suggesting recolonization from two distinct routes forming a secondary contact zone. In the current study, we observe the same change in genetic background between north and south for all three species. However, from genomic data alone, it is difficult to conclude recolonization from both northern and southern routes, especially in silver birch and Scots pine. Nevertheless, as noted in the Introduction, pollen fossil maps support two entry points to Scandinavia for all three species (Brewer et al., 2017), and our genomic analyses do not contradict this scenario.

We also did not attempt to resolve the precise genetic origins of the populations colonizing Sweden from the south and north, as this would require denser sampling across central, northern, and southern Europe. However, a previous nuclear genomic study in *P. abies* indicated that the two recolonization routes involved populations with distinct genetic backgrounds (Zhou et al., 2024). Similarly, mitochondrial data for *P. sylvestris* suggested dual migration into Sweden from two genetically differentiated populations (Sinclair et al., 1999), although Pyhäjärvi et al. (2008) pointed out that alternative explanations for this pattern are possible. For *B. pendula*, a chloroplast DNA (cpDNA) study discussed the possibility of recolonization from two refugia with different genetic compositions (Palmé et al., 2003). However, three birch species are present in Sweden, with hybridization among them, as indicated by extensive and geographically structured chloroplast sharing (Palme et al., 2004), which likely complicates conclusions drawn from cpDNA data.

In general, most forest tree species exhibit F_ST_ values lower than 0.1 (Slavov et al., 2004), and our estimates are consistent with this and previous studies in these three species (Tsuda et al., 2017; Li et al., 2022; James et al., 2023; Bruxaux et al., 2024; Milesi et al., 2024). Despite this overall low differentiation, the limits of the contact zone and the degree of population structure varied among species, likely reflecting differences in migration rates, ecology, and recolonization history. As evidenced by the isolation-by-distance (IBD) patterns and neighborhood size estimates, *P. abies* exhibited the most pronounced structure, with reduced genetic exchange in the northern populations and the contact zone. *B. pendula* showed an intermediate pattern, with gene flow primarily constrained to the contact zone. In line with previous studies (Pyhäjärvi et al., 2019 and references therein; Bruxaux et al., 2024, Kastally et al., 2025), *P. sylvestris* displayed the weakest population structure, characterized by relatively homogeneous migration rates across its range. This confirms that dispersal ability is highest in *P. sylvestris*, intermediate in *B. pendula*, and lowest in *P. abies*, as previously observed over a larger part of their range (Milesi et al., 2024). Additionally, species ecology may also offer an explanation. Birch is a pioneer species, *P. sylvestris* an early colonizer, and *P. abies* typically begins in the understory of already established *Pinus–Betula–Alnus* forests (Seppä et al., 2009). Finally, recolonization history differs between species, which, in part, can explain why the contact zone in *P. abies* is currently more pronounced than in the other two species. Firstly, *P. abies* represents the most recent and constrained spread of a major tree species in northern Europe (Seppä et al., 2009). Secondly, out of the three species, it is only shown for *P. abies* that the trees that entered from north and south likely came from different source populations, as indicated above, having already accumulated substantial divergence (Zhou et al., 2024).

### 2. Cline positions align with environmental gradients

We detected putative signatures of local adaptation in all three species, and a closer examination of the spatial patterns in allele frequencies of candidate SNPs provided further insights into their similarities and differences. Candidate SNPs from selection scans were enriched for clinal alleles in all three species. In *P. abies* and *B. pendula*, nearly all top candidate SNPs followed a continuous gradient, whereas clinal signals in *P. sylvestris* were more variable and less prevalent. The relatively greater variation in model fit and cline parameters for *P. sylvestris* may result from both technical and biological factors: sparse sampling in the central range (60–63° N) can bias cline-parameter estimates (Dufková et al., 2011; Polechová & Barton, 2011) and higher gene flow accelerates cline decay (Figure S10C).

If environmental selection is the primary driver of clinal patterns, allelic cline centers are expected to align with environmental gradient centers. This pattern was clear in *P. abies* and *B. pendula*, with allelic cline centers coinciding with the contact zone and transitions in temperature-related climatic factors. In *P. sylvestris,* no single center dominated, but the top five candidate SNPs had cline centers overlapping with the contact zone and some bioclimatic variables.

Interestingly, *P. abies* also displayed a second cline center for some candidate loci, approximately 350 km from the northernmost part of Sweden—distinct from the centers of the main bioclimatic variables at 600–1000 km (Figure S11). Several non-mutually exclusive explanations could account for this pattern. First, clines may also be shaped by endogenous selection arising from fitness interactions among genes (underdominance or epistasis). Populations with such intrinsic barriers form tension zones when they meet (Bazykin, 1969; Slatkin, 1973). Because tension zones are not fixed by the environment, they can shift geographically; as they move, they may become trapped by exogenous clines maintained by environmental selection, producing coincident clines (Barton, 1979; Bierne et al., 2011). If exogenous selection is weak at the loci under study and gene flow is asymmetric, an endogenous center can shift toward regions of lower dispersal (Barton, 1979). Consistent with this view, the low-migration area revealed by FEEMS in *P. abies* (Figure 2) overlaps the northern cline center (∼350 km), suggesting movement toward a low-density trough. Second, Chen et al. (2019) and Zhou et al. (2024) showed that these far north populations display recent introgression from *Picea obovata* (Siberian spruce) trees, which may be affecting adaptive patterns in this region. These explanations suggest a possible contribution of another form of selection, alongside environmental selection, in maintaining clines at these loci, which might also explain why genome scans identify additional candidate genes beyond those found by GEA.

Overall, we provide evidence for the role of spatially varying environmental selection in shaping clines in the three studied species, in line with previous provenance experiments for these species providing independent evidence for local adaptation (Savolainen et al., 2007; Lascoux et al., 2016). While we speculate that endogenous selection may contribute to heterogeneity in cline centers in *P. abies*, we do not suggest its complete absence in the other two species. The primary driver of clines may be endogenous or exogenous selection, but it is unlikely that they act in complete isolation. Barriers initiated by environmental selection can be reinforced by epistasis, and similarly intrinsic incompatibilities formed in isolation may later couple with environmental selection once gene flow is reestablished (Bierne et al., 2011 and references therein).

### 3. Genomic signals of local adaptation under ongoing gene flow

All three species show genomic signatures of local adaptation along environmental gradients, especially temperature-related factors. Their ability to maintain local adaptation despite high gene flow likely stems from their broad geographic ranges. Local adaptation is easier to achieve over broad spatial ranges than in small, isolated habitats, where gene flow is more likely to be disruptive (Slatkin, 1973). Gene flow can increase genetic variance and reduce drift (Polechová et al., 2009; Polechová & Barton, 2015), enabling populations to respond more effectively to spatial environmental gradients, reducing lag load (Pease et al., 1989), instead of suffering from migration load (Wright, 1984). These conditions are known to be typical of forest trees, which often have broad ranges, high gene flow, and polygenic traits shaped by small-effect loci (Savolainen et al., 2007). Our results are consistent with these previous findings.

Despite sharing a common contact zone and responding to similar environmental gradients, the genomic signatures of local adaptation differed between the three species. We believe that some of the differences among the three species might reflect their relative levels of gene flow. Varying levels of gene flow across such broad ranges, under a polygenic architecture, influence the expected genomic patterns that maintain local adaptation (Tigano & Friesen, 2016) and the species in this study illustrate these predicted patterns. When gene flow is relatively low, selection can counteract migration, enabling adaptive alleles to persist across the genome and producing a diffuse architecture (Yeaman, 2022). The relatively low gene flow in *P. abies*, coupled with the genome-wide distribution of its adaptive alleles, exemplifies a diffuse architecture.

As gene flow increases, large-effect alleles or tightly linked loci may be required to maintain locally adaptive variation, resulting in a concentrated architecture (Bürger & Akerman, 2011; Yeaman & Whitlock, 2011; Yeaman & Otto, 2011). In *B. pendula*, which experiences intermediate gene flow, adaptation is more concentrated. A putative inversion on chromosome 1 potentially captures some adaptive alleles and provides strong resistance against gene flow. A clinal frequency pattern of the inversion along an environmental gradient, combined with relatively homogenized allele frequencies across the rest of the genome, is indicative of spatially varying selection maintaining the inversion (Berry & Kreitman, 1993; Schaeffer, 2008; Cheng et al., 2012). Reduced recombination and migration within this region likely further limit migration load, helping to maintain locally adaptive variants (Berdan et al., 2023). Theory predicts that inversions or tightly linked regions can be advantageous under spatially varying selection by capturing locally adapted alleles (Kirkpatrick & Barton, 2006), including those with positive epistasis (Charlesworth, 1974) or containing fewer deleterious mutations (Nei et al., 1967). Regardless of the specific selective pressures on a newly arisen inversion, empirical studies in various taxa have demonstrated the role of inversions in local adaptation (e.g., Jones et al., 2012; Kapun & Flatt, 2019; Pyhäjärvi et al., 2013; see also Wellenreuther & Bernatchez, 2018, and references therein).

Under high gene flow, adaptive divergence can occur through small shifts in allele frequencies at many loci without strong allelic differentiation, creating a transient adaptive architecture (Le Corre & Kremer, 2012; Yeaman, 2015). This results in a mismatch between phenotypic and genotypic differentiation, where phenotypic divergence is primarily driven by covariances among alleles (Latta, 1998, 2004; Le Corre & Kremer, 2003; Kremer & Le Corre, 2012). The patterns in *P. sylvestris* provide an example for such a transient adaptive architecture.Due to the high level of gene flow, allelic differentiation between populations was very low; however, adaptive signals were present in different parts of the genome, with fewer outliers and less obvious peaks. Previous studies have also reported weak genomic signals of selection (Tyrmi et al., 2020; Bruxaux et al., 2024), despite pronounced phenotypic divergence along latitudinal and longitudinal clines (Notivol et al., 2007; Hall et al., 2021). If a polygenic trait has sufficient genetic variation — which is the case for *P. sylvestris* (Savolainen et al., 2004; Notivol et al., 2007) — local adaptation can draw on various combinations of several small-effect alleles. Despite being prone to swamping, these alleles can temporarily contribute to local adaptation before being lost (Yeaman, 2015). It is also worth mentioning that, despite observing lower and weaker signals in Scots pine compared to silver birch and Norway spruce, we identified more putatively adaptive SNPs than in the previous genomic studies (Bruxaux et al., 2024, Hall et al. 2021). This may be partly due to our use of a more recent and closely related reference genome. Additionally, while Bruxaux et al. (2024) analyzed populations spanning the entire geographic range of *P. sylvestris*, our study focused on a narrower spatial scale within Sweden, which may have increased our power to detect locally adaptive variants.

In addition to the distribution of alleles contributing to adaptation across the genome, we also observed differences in the relative selection strengths of these alleles among species. Consistent with the expectation that local adaptation under increasing gene flow tends to favour stronger selection (Bürger & Akerman, 2011; Yeaman & Whitlock, 2011), selection coefficients for alleles best described by cline Model I increased from *P. abies* to *P. sylvestris* (Figure 7). For alleles best described by cline Model II, the relationship between *P. abies* and *B. pendula* was reversed; however, the difference was smaller and less pronounced than in *P. sylvestris*, where gene flow is strongest. Despite these relative differences, the median selection coefficients across species were weak (approximately 10⁻⁵ to 10⁻²), which is consistent with previous studies in trees suggesting a polygenic basis for local adaptation and the predominant role of small-effect loci (Savolainen et al., 2007).

Transitions between these genetic architectures are, of course, gradual and influenced by the migration–selection balance, which may vary across the genome. These classifications offer a useful framework for interpreting the prominent patterns observed in our data. However, we cannot exclude the potential role of highly linked genomic regions in *P. sylvestris* and *P. abies*, since our dataset represents relatively sparse variant sampling compared to whole-genome sequencing. Indeed, Tyrmi et al. (2020) identified a possible inversion in *P. sylvestris* populations sampled from Finland and Poland. Nevertheless, the relative comparison between species regarding the dominant pattern should remain valid.

### 4. Differences and similarities in adaptive loci

Natural populations in Fennoscandia of all studied species are highly adapted to local conditions, showing geographical cline in growth and phenological traits such as bud set and burst from north to south (Alberto et al., 2013; Chen et al., 2012; Lagercrantz & Ryman, 1990; Milesi et al., 2019; Oksanen, 2021; Savolainen et al., 2007, 2013). However, there is less evidence regarding genetic differentiation along the environmental gradients. Our GEA analyses revealed that a substantial proportion of candidate genes across all species were strongly associated with temperature-related variables (Table S1), suggesting that climate plays a key role in local adaptation. Accordingly, candidate genes for all species were enriched for functions related to climatic stress.

In *P. abies*, as reported by Li et al. (2022), we detected more temperature-related candidate genes than precipitation-related ones. We did not identify the same genes, but our candidate genes were also enriched for the same biological processes, including photoperiodism, flowering, growth, and circadian clock regulation. At the phenotypic level, environmental variables have previously been shown to strongly influence variation in growth and phenology across the natural range of *P. abies* (Lagercrantz & Ryman, 1990; Milesi et al., 2019; Z.-Q. Chen et al., 2021). Analyses of long-term forest experiments assessing growth response to climatic factors in Sweden have also shown that *P. abies* is more sensitive to early summer precipitation than *P. sylvestris*, with radial growth particularly affected at lower latitudes (Ogana et al., 2024).

In *B. pendula*, a limited number of enriched GO terms were detected, which may be due to the targeted sequencing strategy focusing on a few exonic regions. However, enrichment for biological processes such as primary shoot apical meristem specification is linked to growth and bud break, traits for which different experiments have shown genotypic variation in response to increased temperature (reviewed in Oksanen, 2021). Additionally, Salojärvi et al. (2017) identified several *B. pendula* candidate genes displaying clinal variation in relation to environmental variables, including *MED5A* (encoding Mediator complex protein MED5A). Consistent with that study, we also identified *MED5A* among our candidate genes, which is associated with growth in *Arabidopsis*.

Enrichment analyses were also limited for *P. sylvestris* and most enriched candidate genes were related to intra-cellular biological processes. However, an enriched GO term related to the gene SSR1 was linked to the rhythmic biological process. This gene is important for photoperiodic and photoperiod-independent regulation of flowering in *A. thaliana* and its involvement in adaptation along latitude has been shown before in *P. abies* (Li et al. 2022). Phenotypic studies including transfers) and common-garden experiments (Hall et al., 2021) have revealed a variation in frost tolerance in Swedish populations of *P. sylvestris,* although no strong genetic signal has been detected for this trait. Similarly, we could not associate our outliers with frost tolerance.

Overlap in candidate genes and GO terms across species was lower than expected by chance. This low functional overlap may partly reflect technical limitations rather than true biological differences. Exome sequencing in *P. abies* and *B. pendula* used different targeting strategies: the former covered most genes in the genome, while the latter targeted a reduced gene set, similar to the GBS approach used for *P. sylvestris*. In addition, the gene annotation of the closest reference genome for *P. sylvestris* is less complete, further limiting functional interpretation. Nevertheless, the detection of adaptive loci in *P. sylvestris* despite these limitations supports the idea of highly polygenic adaptation, potentially involving regulatory regions. Similarly, across all species, the polygenic nature of adaptive traits—where different genes may contribute to similar phenotypes—may also explain the low overlap.

In summary, our comparative analysis of three boreal tree species reveals a shared secondary contact zone in central Sweden that separates two main genetic clusters and aligns with a major climatic transition. In all three species, this contact zone appears to be maintained by spatially varying environmental selection, as evidenced by clinal patterns in allele frequencies of candidate loci along temperature-related environmental gradients. The differences among species in the limits of the contact zone and in the degree of genetic differentiation across it reflect variation in gene flow and the resulting population structure. These differences in gene flow levels are also reflected in the species-specific adaptive architectures, providing empirical support for theoretical expectations about local adaptation under migration. Functional enrichment of genes involved in growth, phenological timing, and environmental stress responses further supports climate as the main driver of polygenic local adaptation in all three species. Altogether, our study highlights the value of comparative contact zone analyses for understanding how gene flow and selection jointly shape genomic variation. More specifically, it shows the importance of investigating local adaptation in forest tree species for predicting how natural populations may respond to ongoing climate change and for informing breeding strategies based on genomic prediction.

## Acknowledgements

The present project has been funded by the support of the Swedish Energy Agency through the Trees for Me Center, as well as by the Nilsson Ehle Endowment from the Swedish Royal Physiographic Society of Lund and the Swedish Phytogeographical Society. We thank Tanja Pyhäjärvi and Parsi Rastas for access to the improved assembly and annotation of *Pinus tabuliformis*, Janek Sendrowski for providing us with the initial data of *Betula pendula* used for this project, and Rachid Cheddadi, Heikki Seppä and Thomas Giesecke for help with understanding the pollen fossil record. The computations and data storage were enabled by resources under the project NAISS 2024/22-11 (UPPMAX 2025/2-229), provided by the Swedish National Infrastructure for Computing (SNIC) at UPPMAX.

## Data Availability

The data and codes necessary to replicate the analyses presented in the manuscript are available at: https://github.com/mpilarhego/SWEtrees_comp.The raw-reads for the *B. pendula* samples are available in ENA under project number PRJEB96483.

## Author Contributions

PM, ML and BY conceived and designed the study. PHE, BY, and PM devised the analysis strategy and performed all data analyses. QZ and LL contributed data for *P. abies* and *B. pendula*, respectively. PHE, BY, ML, and PM wrote the manuscript. ML and PM supervised the study and ML acquired funding.

