## Supplementary_Figures_Tables for "When South meets North: a joint contact zone coinciding with environmental gradients in three boreal tree species": SWETrees_SuppFiguresTables_revised_smaller.pdf

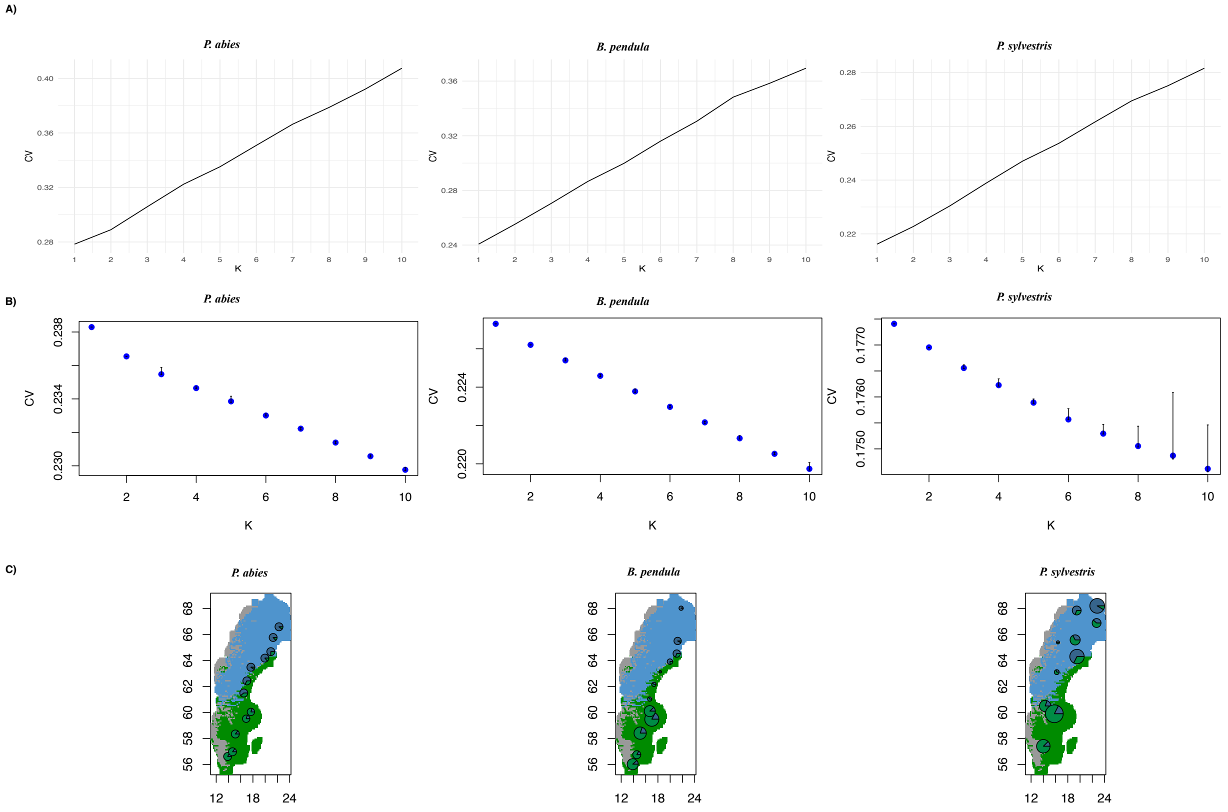


**Figure S1:** Cross-validation errors for different K values from A) ADMIXTURE and B) TESS3 analysis. C) Ancestry proportions for each population based on the TESS3 (K = 2), with colors representing different ancestry components. The background colors corresponds to the climatic zones, climatic zone 1 (blue), climatic zone 2 (green), climatic zone 3 (grey).

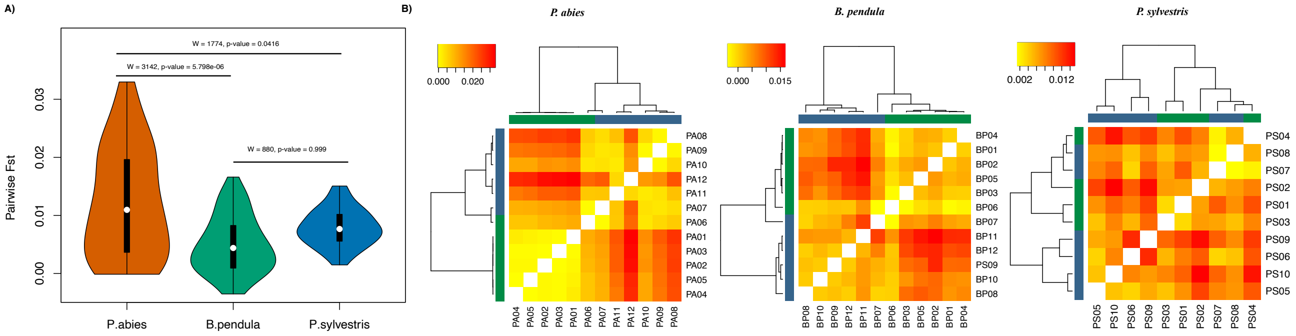


**Figure S2:** A) Distribution of Pairwise $F_{ST}$ values for each species, along with one-sided Wilcoxon rank-sum test results between species B) Pairwise $F_{ST}$ heatmap and dendrogram. The blue and green colors in the rows and columns correspond to the northern and southern genetic cluster information of the populations, respectively.

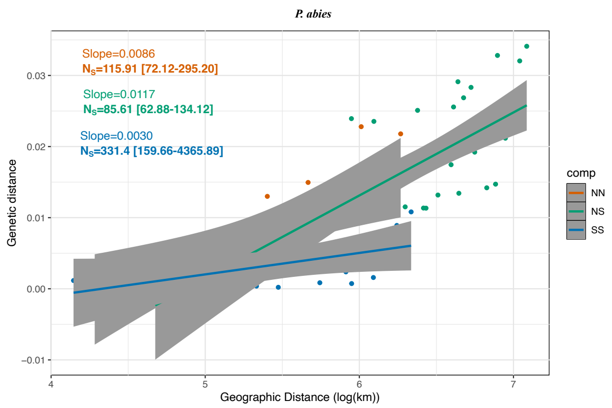


**Figure S3:** Isolation by distance (IBD) patterns calculated separately for northern-northern (NN), northern-southern (NS), or southern-southern (SS) populations of Norway spruce (*P. abies*), along with the neighborhood size ($N_S$) estimates.

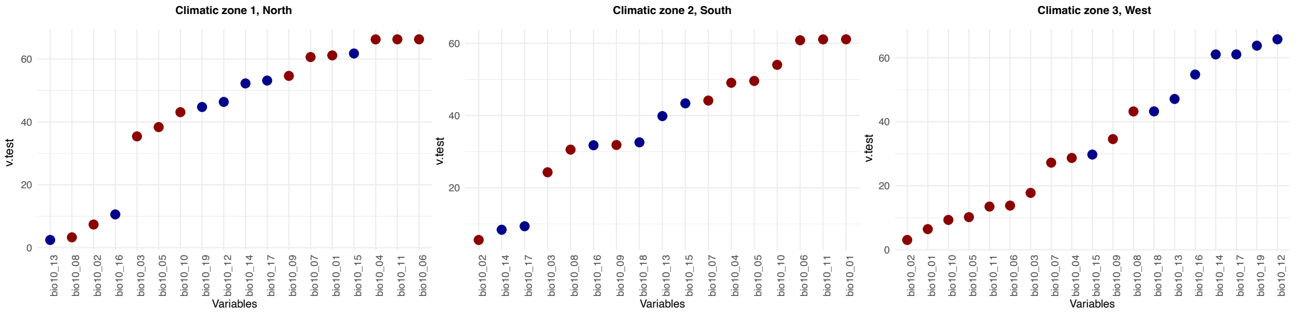


**Figure S4:** Contribution of climatic variables to each environmental cluster. The contribution is quantified by the v-test statistic, with higher values indicating a stronger influence. Red dots represent temperature related variables, blue dots represent precipitation related variables. The three most influential variables per cluster are: Cluster 1 — Temperature seasonality (bio4), Minimum temperature of the coldest month (bio6), and Mean temperature of the coldest quarter (bio11); Cluster 2 — Annual mean temperature (bio1), Minimum temperature of the coldest month (bio6), and Mean temperature of the coldest quarter (bio11); Cluster 3 — Annual precipitation (bio12), Precipitation of the driest quarter (bio17), and Precipitation of the coldest quarter (bio19). Definitions of all variables are provided in Table S1.

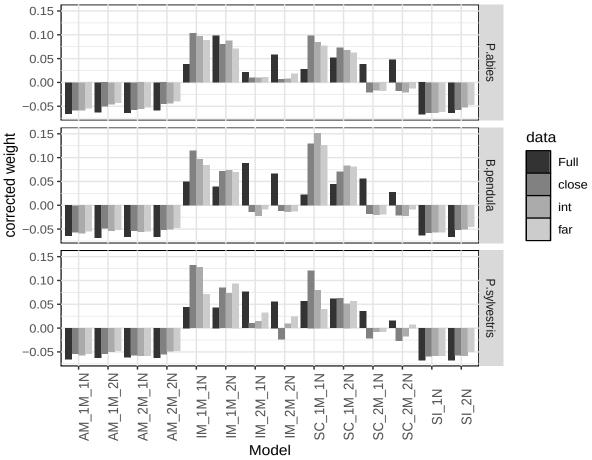


**Figure S5:** Model weight of 14 models (adjusted for the uniform distribution model weight) for each species and dataset; full or separated into three subsets based on the genetic distance between individuals (close, intermediate, and far).

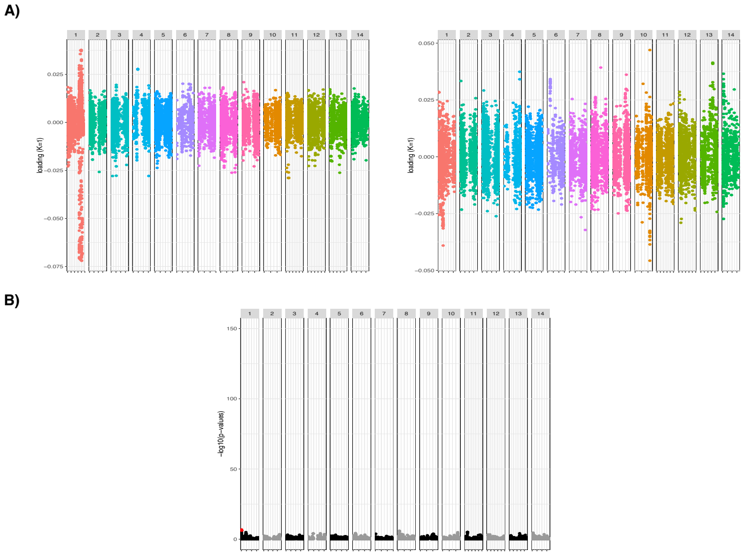


**Figure S6:** Loadings before and after filtering a region in chromosome 1 (~9Mbp), and a genome scan post-filtering for *B. pendula*. A) Loadings showing high correlation between each genotype p-values (*pcadapt*) and K=1 principal component before (left panel) and after (right panel), excluding the genomic region (Chr1) of high contribution of outliers. B) Manhattan plot of genome scan analyses (*pcadapt*) after exclusion of ~9 Mbps region in Chromosome 1.

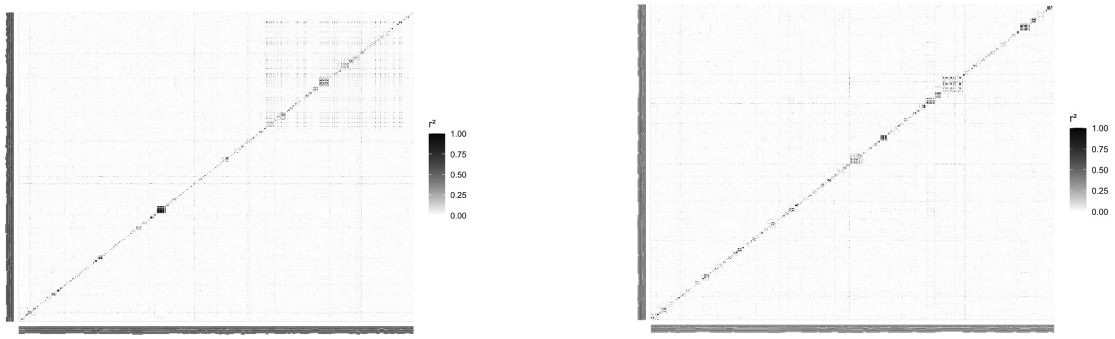


**Figure S7:** Linkage disequilibrium (LD) heatmap with pairwise $r^2$ values calculated between every SNP in chromosome 1 (Left panel) and in chromosome 2 (Right panel) as a comparison.

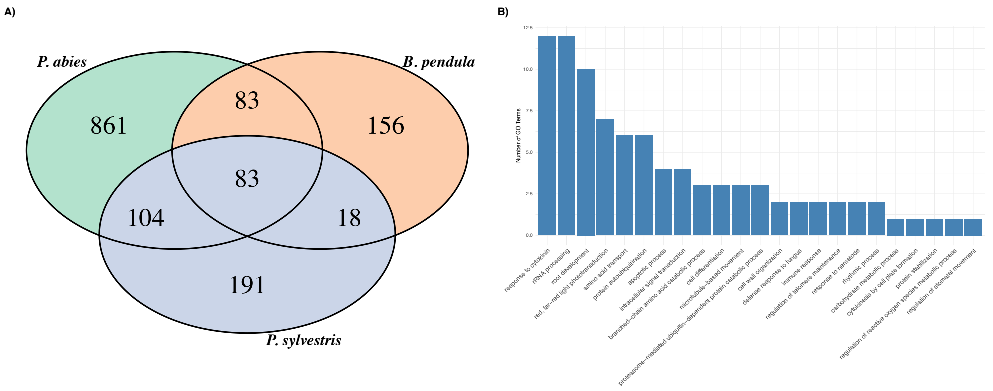


**Figure S8:** A) Unique and overlapped GO terms for candidate genes across the three studied species. B) Functions of 91 common GO terms

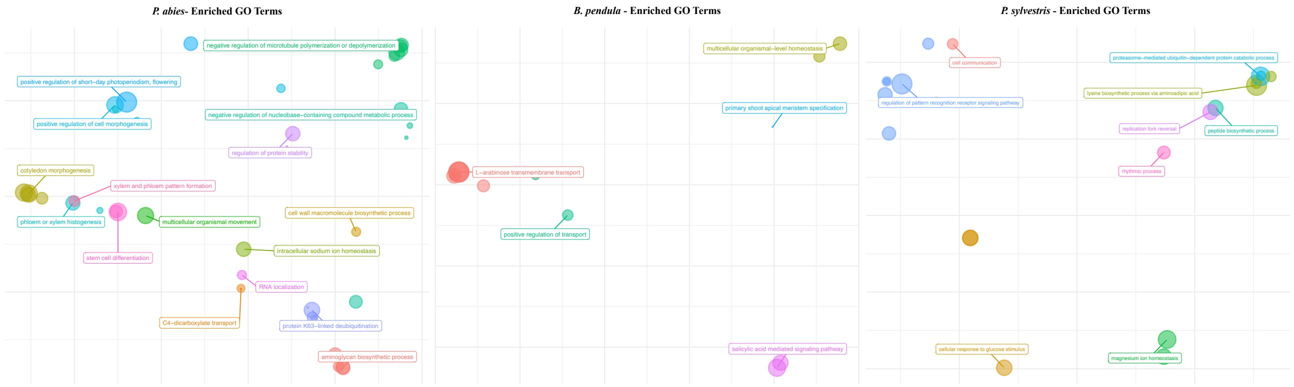


**Figure S9:** Enrichment analyses of GO terms (biological process) for candidate genes of *P. abies, B. pendula* and *P. sylvestris*.

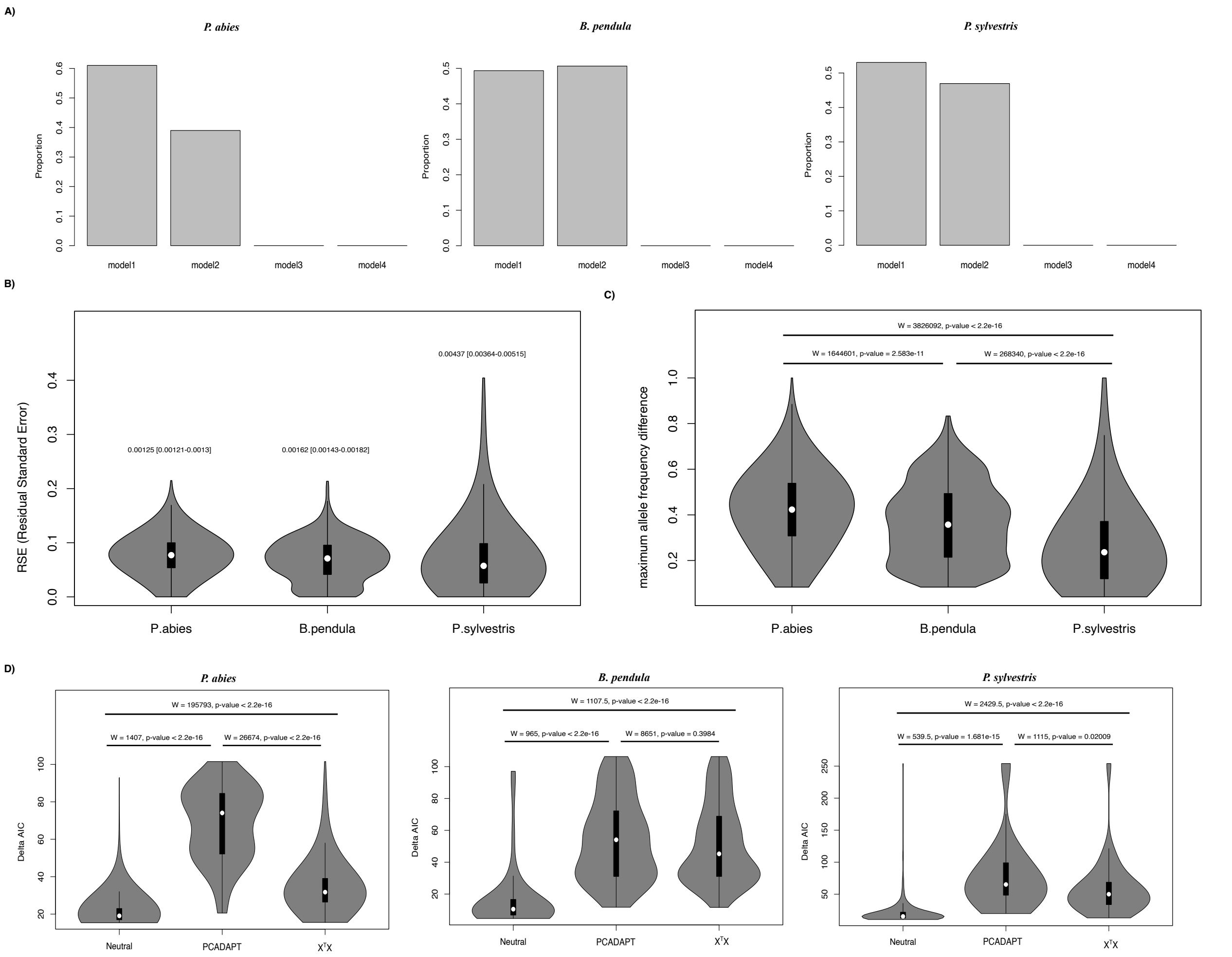


**Figure S10:** A) Proportion of best fitted cline models for clinal alleles of each species. Model I and II represent sigmoid cline, while model III and IV represent stepped cline. All clinal alleles followed a sigmoid shape. B) Distribution of residual standard errors (RSE) for all clinal alleles in each species. The variation in distributions differs significantly. The values above the distributions indicate variance values and confidence intervals.. C) Distribution of maximum allele frequency differences between populations for all clinal alleles in each species, along with results of the Wilcoxon rank-sum test between species. D) Distribution of AIC differences between the null model and cline models for different sets of clinal alleles (Neutral, X$^T$X outliers, and *pcadapt* outliers). The values indicate the results of the Wilcoxon rank-sum test.

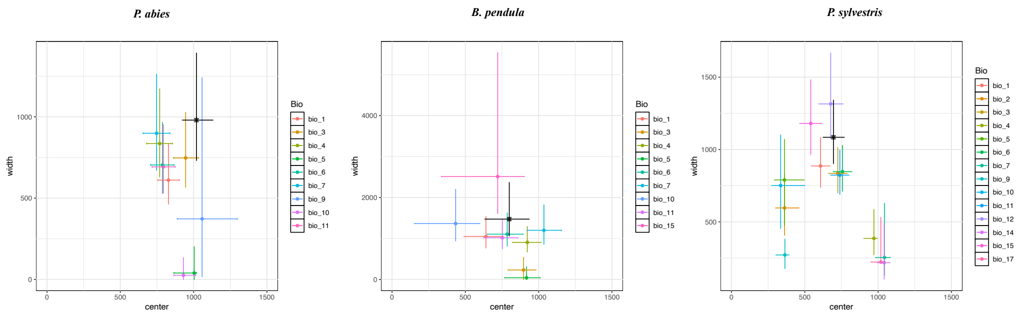


**Figure S11:** Cline parameters (with 95% confidence intervals) of environmental values for variables showing significant cline patterns in each species. Black dot shows the cline parameters for the first principal component (PC1) of the principal component analysis conducted on the 19 bioclimatic variables. Environmental and PC1 values are taken for locations of each population.

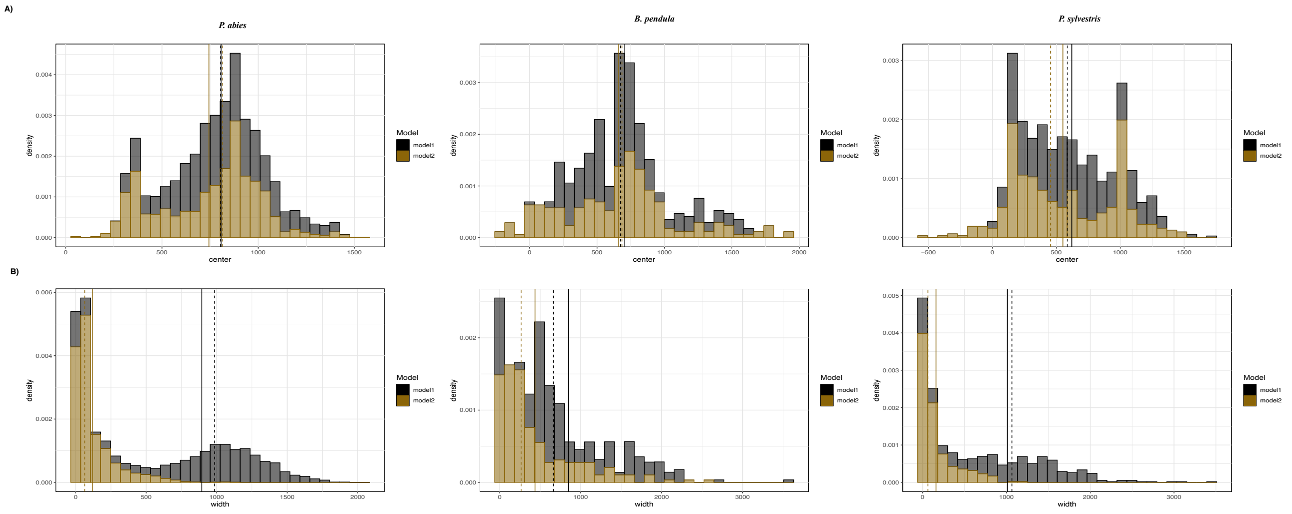


**Figure S12:** A) Center parameter distribution separately for model I and II. B) Width parameter distribution separately for model I and II. Dashed and solid lines correspond to median and mean values, respectively, for each model.

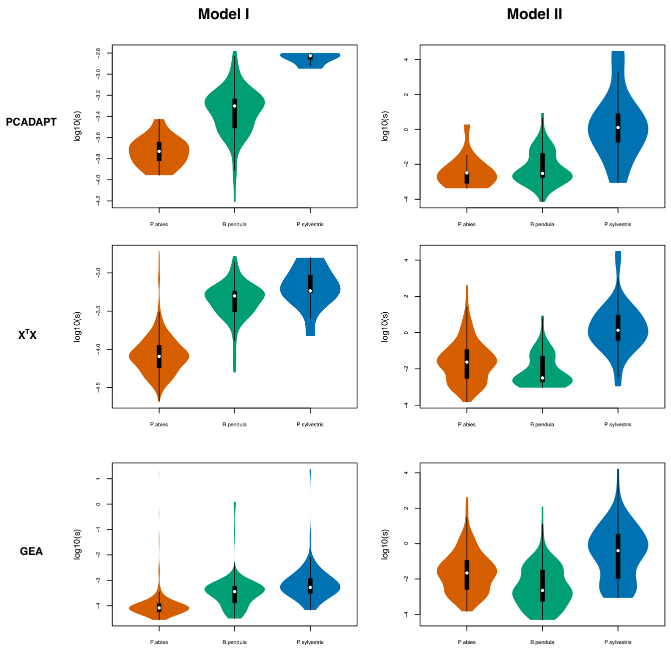


**Figure S13:** Selection coefficients calculated from cline width parameters separately for each selection scan and cline model.

### SUPPLEMENTARY TABLES

**Table S1:** Summary of candidate SNPs and genes identified by genome scans and GEA analyses across all species. For total number of SNPs and genes, proportion compared to full data is given in brackets.

| Analyses | *P. abies* | | *B. pendula* | | *P. sylvestris* | |
|---|---|---|---|---|---|---|
| | SNPs | genes | SNPs | genes | SNPs | genes |
| *pcadapt* | 39 | 17 | 119 | 46 | 42 | 21 |
| $X^TX$ | 806 | 363 | 137 | 51 | 92 | 43 |
| Annual Mean Temperature (Bio1) | 68 | 32 | 121 | 48 | 50 | 29 |
| Mean Diurnal Range (Bio2) | 14 | 10 | 29 | 16 | 33 | 21 |
| Isothermality (Bio3) | 35 | 27 | 96 | 42 | 43 | 23 |
| Temperature Seasonality (Bio4) | 62 | 32 | 86 | 43 | 29 | 18 |
| Max Temperature of Warmest Month (Bio5) | 24 | 18 | 71 | 33 | 37 | 25 |
| Min Temperature of Coldest Month (Bio6) | 75 | 36 | 99 | 45 | 39 | 22 |
| Temperature Annual Range (Bio7) | 48 | 24 | 32 | 22 | 25 | 14 |
| Mean Temperature of Wettest Quarter (Bio8) | 12 | 10 | 0 | 0 | 13 | 8 |
| Mean Temperature of Driest Quarter (Bio9) | 26 | 17 | 8 | 8 | 25 | 8 |
| Mean Temperature of Warmest Quarter (Bio10) | 27 | 19 | 80 | 37 | 35 | 24 |
| Mean Temperature of Coldest Quarter (Bio11) | 80 | 36 | 113 | 46 | 39 | 22 |
| Annual Precipitation (Bio12) | 13 | 9 | 2 | 2 | 38 | 21 |
| Precipitation of Wettest Month (Bio13) | 16 | 12 | 2 | 2 | 2 | 1 |
| Precipitation of Driest Month (Bio14) | 11 | 8 | 2 | 2 | 25 | 13 |
| Precipitation Seasonality (Bio15) | 13 | 10 | 0 | 0 | 46 | 28 |
| Precipitation of Wettest Quarter (Bio16) | 14 | 11 | 3 | 3 | 13 | 5 |
| Precipitation of Driest Quarter (Bio17) | 10 | 6 | 2 | 2 | 31 | 11 |
| Precipitation of Warmest Quarter (Bio18) | 18 | 13 | 1 | 1 | 10 | 3 |
| Precipitation of Coldest Quarter (Bio19) | 5 | 4 | 2 | 2 | 36 | 18 |
| Unique Total | 999 (0.17%) | 495 (1.77%) | 163 (0.31%) | 66 (3.05%) | 197 (0.22%) | 96 (1.65%) |
| Unique Total GEA (all Bio) | 300 (0.05%) | 183 (0.65%) | 142 (0.27%) | 61 (2.82%) | 142 (0.16%) | 76 (1.30%) |

**Table S2:** Coefficients of overlap for candidate genes from the different genome scans and GEA analyses, calculated as the intersect across datasets divided by the minimum dataset size. P: *pcadapt*, X: $X^TX$, G: GEA.

| Overlap Coefficient | *P. abies* | *B. pendula* | *P. sylvestris* |
|---|---|---|---|
| **P∩X∩G** | 0.76 | 0.93 | 0.57 |
| **P∩X** | 1.00 | 0.98 | 0.86 |
| **X∩G** | 0.28 | 0.90 | 0.56 |
| **P∩G** | 0.76 | 0.96 | 0.67 |

**Table S3:** Coefficients of overlap of GO terms for candidate genes and background genes (from SNPs in initial VCF file) in all species and pairwise comparisons between them (df =1, *P*-values < 1.3e-51).

| **Comparison** | **GO terms** | **Common** | **Total** | **Ratio (95% CI)** | $\chi^2$ |
|---|---|---|---|---|---|
| All species | candidate | 83 | 1496 | 0.06 (0.04,0.07) | 1181 |
| | all | 2773 | 7919 | 0.35 (0.34,0.36) | 710 |
| Pabies_Bpendula | candidate | 166 | 1305 | 0.13 (0.11,0.15) | 724 |
| | all | 3219 | 7638 | 0.42 (0.41,0.43) | 188 |
| Bpendula_Psylvestris | candidate | 101 | 635 | 0.16 (0.13,0.19) | 294 |
| | all | 2810 | 5936 | 0.47 (0.46,0.49) | 17 |
| Pabies_Psylvestris | candidate | 187 | 1340 | 0.14 (0.12,0.16) | 695 |
| | all | 5071 | 7818 | 0.65 (0.64,0.66) | 690 |

**Table S4:** Proportion of clinal alleles in different allele sets (neutral and selection outliers) and the adjusted *P*-values (Bonferroni) from hypergeometric test for enrichment compared to all dataset are given inside brackets. Significantly enriched datasets are depicted in bold.

| **Allele Set** | ***P. abies*** | ***B. pendula*** | ***P. sylvestris*** |
|---|---|---|---|
| Neutral | 0.0105 (0.099) | 0.0096 (1.000) | **0.0134 (1.36e-11)** |
| *pcadapt* | **1.000 (4.80e-21)** | **1.000 (1.06e-177)** | **0.692 (2.54e-23)** |
| $X^TX$ | **0.932 (0.00e+0)** | **1.000 (1.45e-206)** | **0.724 (1.37e-51)** |
| Annual Mean Temperature (Bio1) | **0.985 (2.51e-76)** | **1.000 (7.00e-181)** | **0.940 (4.89e-64)** |
| Mean Diurnal Range (Bio2) | 0.214 (3.50e-01) | **1.000 (6.38e-27)** | **0.879 (3.95e-39)** |
| Isothermality (Bio3) | **0.743 (4.06e-25)** | **0.989 (4.10e-138)** | **0.930 (2.60e-54)** |
| Temperature Seasonality (Bio4) | **0.903 (8.43e-50)** | **1.000 (3.25e-126)** | **1.000 (1.28e-35)** |
| Max Temperature of Warmest Month (Bio5) | **0.500 (1.55e-11)** | **0.986 (1.20e-95)** | **0.919 (7.48e-46)** |
| Min Temperature of Coldest Month (Bio6) | **0.933 (5.80e-66)** | **1.000 (2.74e-146)** | **1.000 (1.02e-51)** |
| Temperature Annual Range (Bio7) | **0.854 (7.66e-46)** | **1.000 (6.79e-30)** | **0.960 (6.04e-28)** |
| Mean Temperature of Wettest Quarter (Bio8) | 0.000 (1.000) | 0.000 (1.000) | **0.769 (5.66e-10)** |
| Mean Temperature of Driest Quarter (Bio9) | **0.269 (8.40e-05)** | **1.000 (3.55e-06)** | **0.840 (5.08e-26)** |
| Mean Temperature of Warmest Quarter (Bio10) | **0.407 (1.72e-08)** | **0.975 (1.11e-105)** | **0.914 (1.97e-42)** |
| Mean Temperature of Coldest Quarter (Bio11) | **0.975 (1.17e-87)** | **1.000 (3.30e-168)** | **1.000 (1.03e-51)** |
| Annual Precipitation (Bio12) | 0.000 (1.000) | 0.000 (1.000) | **0.815 (3.00e-36)** |
| Precipitation of Wettest Month (Bio13) | 0.000 (1.000) | 0.000 (1.000) | 0.00 (1.00) |
| Precipitation of Driest Month (Bio14) | 0.000 (1.000) | 0.000 (1.000) | **0.760 (1.20e-18)** |
| Precipitation Seasonality (Bio15) | 0.154 (1.000) | 0.000 (1.000) | **0.934 (2.12e-57)** |
| Precipitation of Wettest Quarter (Bio16) | 0.000 (1.000) | 0.000 (1.000) | **0.466 (8.54e-03)** |
| Precipitation of Driest Quarter (Bio17) | 0.000 (1.000) | 0.000 (1.000) | **0.806 (2.05e-24)** |
| Precipitation of Warmest Quarter (Bio18) | 0.000 (1.000) | 0.000 (1.000) | **0.600 (7.24e-03)** |
| Precipitation of Coldest Quarter (Bio19) | 0.000 (1.000) | 0.000 (1.000) | **0.805 (3.06e-26)** |
